# Radiation of nitrogen-metabolizing enzymes across the tree of life tracks environmental transitions in Earth history

**DOI:** 10.1101/2020.05.01.064543

**Authors:** Chris Parsons, Eva Stüeken, Caleb Rosen, Katherine Mateos, Rika Anderson

## Abstract

Nitrogen is an essential element to life and exerts a strong control on global biological productivity. The rise and spread of nitrogen-utilizing microbial metabolisms profoundly shaped the biosphere on the early Earth. Here we reconciled gene and species trees to identify birth and horizontal gene transfer events for key nitrogen-cycling genes, dated with a time-calibrated tree of life, in order to examine the timing of the proliferation of these metabolisms across the tree of life. Our results provide new insights into the evolution of the early nitrogen cycle that expand on geochemical reconstructions. We observed widespread horizontal gene transfer of molybdenum-based nitrogenase back to the Archean, minor horizontal transfer of genes for nitrate reduction in the Archean, and an increase in the proliferation of genes metabolizing nitrite around the time of the Mesoproterozoic (∼1.5 Ga). The latter coincides with recent geochemical evidence for a mid-Proterozoic rise in oxygen levels. Geochemical evidence of biological nitrate utilization in the Archean and early Proterozoic may reflect at least some contribution of dissimilatory nitrate reduction to ammonium (DNRA) rather than pure denitrification to N_2_. Our results thus help unravel the relative dominance of two metabolic pathways that are not distinguishable with current geochemical tools. Overall, our findings thus provide novel constraints for understanding the evolution of the nitrogen cycle over time and provide insights into the bioavailability of various nitrogen sources in the early Earth with possible implications for the emergence of eukaryotic life.

## Introduction

Nitrogen is a critical element to life on Earth, important as an essential building block in the synthesis of biological molecules, and for its role in redox reactions for microbial energy metabolism. It is often a limiting nutrient in marine and terrestrial environments and likely had a significant influence on the evolutionary trajectory of the biosphere over Earth’s history. The nitrogen cycle is largely controlled by a variety of microorganisms that enzymatically catalyze the reduction and oxidation of nitrogen at various redox states (Figure 1). Reconstructing the genetic proliferation of these enzymes across the tree of life through gene birth, duplication, loss, and horizontal gene transfer can therefore provide novel insights into the evolution of the biosphere and its productivity over time.

**Figure 1.**
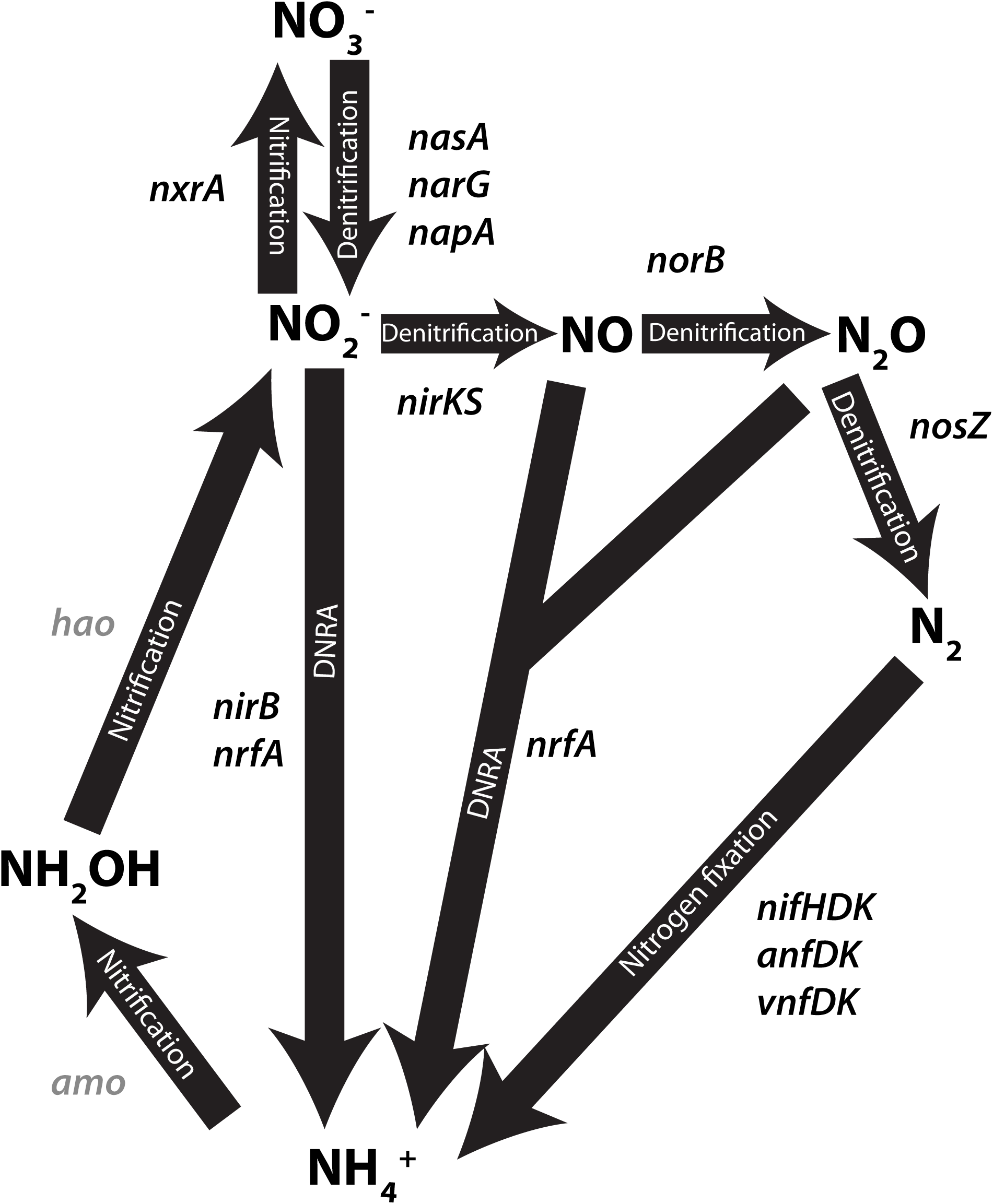
Schematic of the biological nitrogen cycle. Arrows are labelled with the pathway to which they belong. All genes examined in this study are labeled next to the step they catalyze. Genes without sufficient data for subsequent analysis are labeled in grey. Adapted from Canfield et al. (2010).

The most important steps in Earth’s nitrogen cycle are largely catalyzed by microbes, including the first crucial step of reducing molecular nitrogen to bioavailable forms (Zerkle & Mikhail, 2017; Kuypers *et al*., 2018). Nitrogen fixation is catalyzed by nitrogenase, of which there are three varieties, distinguished by the metals in their associated active site cofactors: Nif (Fe-Mo), Vnf (Fe-V), and Anf (Fe-Fe) (Joerger *et al*., 1988; Miller & Eady, 1988) (Figure 1). The ability to fix nitrogen is spread across a wide range of archaeal and bacterial lineages, but does not occur in eukaryotes (Dos Santos *et al*., 2012; Gaby & Buckley, 2014). Importantly, nitrogenase is strongly inhibited by oxygen, forcing nitrogen fixers to develop various means to reduce their intracellular oxygen concentrations or to confine themselves to suboxic environments (Gallon, 1981). Ammonium produced from nitrogen fixation or ammonification is converted to organic forms of fixed nitrogen by a variety of enzymes or, in the presence of oxygen, oxidized to nitrite (NO_2_^-^) or nitrate (NO_3_^-^) through the chemoautotrophic nitrification pathway via the enzymes Amo and Hao (Figure 1). In environments with insufficient O_2_ concentrations to support aerobic respiration, nitrate and nitrite can be utilized as alternative terminal electron acceptors through the denitrification pathway; consequently, denitrification rates are highest in suboxic conditions, including swamps and marine oxygen minimum zones (Canfield *et al*., 2005; Löscher *et al*., 2012; Voss *et al*., 2013). Reduction of nitrate to nitrite, nitric oxide (NO), nitrous oxide (N_2_O) and dinitrogen (N_2_) is catalyzed by the Nas, Nar, Nap, Nir, Nor, and Nos enzymes, respectively. Dissimilatory nitrate reduction to ammonium (DNRA) also reduces nitrate via the enzymes Nar, Nap, Nir and Nrf (Figure 1), but this pathway retains fixed nitrogen as ammonium and may therefore have been a critical metabolism in nutrient-starved ecosystems. DNRA, while less understood than denitrification, has been shown to be a major nitrate sink in a variety of aquatic systems, especially warm intertidal zones (Giblin *et al*., 2013) and it may be dominant under ferruginous conditions, as suggested by modern analogue studies (Michiels *et al*., 2017).

Given the importance of nitrogen as a building block of life, as an energy source for microbes, and as the most abundant element in the Earth’s atmosphere, better constraining the evolutionary history of the nitrogen cycle is important for understanding, among other things, variation in global primary productivity and atmospheric pressure over time. The bioavailability and cycling of important limiting nutrients through Earth’s history, including nitrogen, would have been important for biological productivity and the rise of early eukaryotic algae (Anbar & Knoll, 2002; Sánchez-Baracaldo *et al*., 2014; Isson *et al*., 2018). The relative abundance of nitrogenous gases in the atmosphere could also have had important implications for atmospheric pressure as well as planetary climate during the Archean. Potential changes in atmospheric pressure during the Archean may have resulted from biological N_2_ drawdown (Som *et al*., 2016), whereas the greenhouse gas nitrous oxide (N_2_O), produced as part of the nitrogen cycle via nitrification/denitrification, may have contributed to planetary warming when the Sun was younger and fainter (Buick, 2007; Roberson *et al*., 2011; Stanton *et al*., 2018). Finally, one of the most important questions for the early evolution of life is understanding when fixed nitrogen first became widely available, and through what means. Experimental data suggest that fixed nitrogen can be produced during lightning reactions and under hydrothermal conditions (e.g., Brandes *et al*., 1998; Navarro-González *et al*., 2001), and either or both of these sources were likely pivotal for the origin of life. However, the invention of biological N_2_ fixation would have made Earth’s biosphere less dependent on abiotic reactions and likely spurred primary productivity.

The question of how these metabolisms unfolded over Earth’s history has previously been addressed with both geochemical and phylogenetics-based approaches. Geochemical approaches, relying on the reconstruction of metabolisms based on the rock record, have suggested that biological nitrogen fixation emerged early (Stüeken *et al*., 2016a; Koehler *et al*., 2019; Ossa Ossa *et al*., 2019) and that the nitrogen cycle expanded considerably during the Neoarchean (2.8-2.5 Ga) and Paleoproterozoic (2.5-1.8 Ga) (Garvin *et al*., 2009a; Godfrey & Falkowski, 2009; Zerkle *et al*., 2017a; Kipp *et al*., 2018; Koehler *et al*., 2018; Luo *et al*., 2018). Phylogenetics studies, relying on sequence data, reconstruct the evolutionary history of genes of interest (e.g. Jones *et al*., 2008; Boyd *et al*., 2011b; Garcia *et al*., 2020), and some molecular clock studies have yielded conservative estimates for the approximate timing of an enzyme’s origin (Raymond *et al*., 2004; Boyd *et al*., 2011b; Boyd & Peters, 2013). While each approach provides valuable insights, they both have weaknesses. Geochemical data cannot reliably distinguish between all enzymatic pathways, because the isotopic effects of some reactions (e.g., denitrification, DNRA and ANAMMOX) are too similar to each other. Furthermore, geochemical data, which are typically collected from bulk rock samples, only preserve evidence of the most dominant metabolisms and may therefore not capture the origin of new enzymes until they gain ecological significance. Conversely, the phylogenetic approach of dating the antiquity of enzymes does not show when these enzymes gained ecological dominance. Furthermore, most phylogenetic studies have so far focused on nitrogenase, leaving the evolutionary history of most nitrogen-cycling enzymes poorly constrained. Additional work is therefore needed to address key questions about the dynamics of the nitrogen cycle on the early Earth and its evolution over time. The proliferation of sequencing data over the past decade has made vast amounts of genomic data available, which can provide novel insights into the evolution of nitrogen cycling genes over time.

As a new approach to these questions, we track the timing of birth, speciation, duplication, loss, and horizontal gene transfer events for genes involved in each step of the nitrogen cycle, which can provide contextual information for the rise and spread of key nitrogen-cycling genes across the tree of life. We place particular focus on the acquisition of new genes via horizontal gene transfer (HGT), which is common in microbial lineages and is a crucial evolutionary mechanism by which a clade of organisms can develop new and useful phenotypes without the evolutionary cost associated with independently evolving genes (Beiko *et al*., 2005; Gogarten & Townsend, 2005). Studies of gene gain and loss have revealed a history of widespread HGT throughout the microbial tree of life, which we have attempted to leverage as a means to attribute trends in microbial evolution to events in Earth history (Koonin *et al*., 2001; Mirkin *et al*., 2003). Many of the major genes in the nitrogen cycle have been shown to have experienced extensive HGT, presumably due to their modularity and their strong dependence on oxygen availability (Stolz & Basu, 2002; Kechris *et al*., 2006; Jones *et al*., 2008). We tracked birth, speciation, duplication, loss, and HGT of genes in the nitrogen cycle over time by comparing the phylogenies for specific nitrogen-metabolizing genes to a time-calibrated tree of life to demonstrate when these genes first arose and then spread across the tree of life on the early Earth.

## Materials and Methods

### Genome Selection and Compilation

The construction of both the gene and species trees for this study was based upon the manual curation of a genome database containing 308 genomes (including 254 bacterial and archaeal genomes) that served as the basis for the species tree and was subsequently searched to find genes related to nitrogen metabolism. Assembled genomes were downloaded from ggKBase (Hug *et al*., 2016) and the NCBI assembly database (Kitts *et al*., 2016). Additionally, 6 genomes were collected from a recent study identifying novel nitrogen fixers (Delmont *et al*., 2018). In constructing the tree, we included at least one genome from each bacterial or archaeal phylum represented in the most recent comprehensive tree of life (Hug *et al*., 2016) in order to create a tree fully representative of our current understanding of microbial diversity. It also includes a set of genomes associated with a database of *nifH* genes (Gaby & Buckley, 2014). Some eukaryotic genes were included for construction of the tree, but for this study we focused only on archaeal and bacterial genomes for identification of nitrogen cycling genes. Relative to archaea and bacteria, eukaryotes play a more minor role in the nitrogen cycle—while some species of fungi and protists reduce nitrate or nitrite to more reduced forms of nitrogen, there are no known eukaryotes that mediate nitrogen fixation, nitrification, DNRA, or anammox (Stein & Klotz, 2016).

### Species tree and chronogram construction

To create the species tree, all bacterial and archaeal genomes in our database were mined for single copy ribosomal protein sequences L2, L3, L4, L5, L6, L14, L15, L16, L18, L22, L24, S3, S8, S10, S17, and S19 using Phylosift (Darling *et al*., 2014), with the isolate and best hit command line flags. These sixteen ribosomal proteins represent the same proteins used to create a recent comprehensive tree of life (Hug *et al*., 2016) and, in the case of eukaryotes, ribosomal sequences were directly drawn from their dataset. All genomes included in the dataset contained fewer than 50% gaps in the alignment. These 16 single-copy ribosomal proteins were concatenated to create a final alignment of 2897 characters for phylogenetic reconstruction and molecular clock evaluation.

Alignments for the species trees were made using the Phylosift pipeline (Darling *et al*., 2014) and curated to only include the target ribosomal proteins listed above. The phylogeny was constructed using RAxML v.8.2.9 with 100 rapid bootstraps (Stamatakis, 2014). The CAT model, which calculates site-specific evolutionary rates, was used with an LG substitution matrix to construct the species tree based on reference marker genes (the species tree). The root for the species tree was placed in the Bacterial domain (Fournier & Gogarten, 2010).

### Chronogram Construction

The species tree was converted into chronograms using PhyloBayes (Lartillot *et al*., 2009) using different clock models and calibration points in order to test the sensitivity of our results to variation in Phylobayes parameters. The root age was set via a normally distributed gamma root prior according to the liberal or conservative calibration points set in Table 1 and Supplementary Table 1, with standard deviation set to 200 in accordance with previous studies (Magnabosco *et al*., 2018). We tested two separate sets of calibration points, one liberal (which represents the earliest date for which there is any evidence of a given event based on the current scientific literature) and one conservative (which represents the earliest date for which there is the most consensus for a given event based on the current scientific literature), to test the sensitivity of methodology (Table 1 and Supplementary Table 1). For each set of internal calibration points, the ages in the calibration files were set as the hard lower bound for the analysis. The liberal calibration points shown in Supplementary Table 1 yielded unrealistic root ages (>4.5 Ga) and therefore were not used for further analyses.

**Table 1.** Fossil calibration points used in Phylobayes runs. Calibration points were set as the hard constraint indicating the latest date by which a specific clade split. The selected time points reflect the dates for which there is the most consensus.

| Calibration Event | Date (Mya) | Refs |
| --- | --- | --- |
| LUCA (set as root prior) | 3800, 200 S.D. | Schidlowski *et al.*, 1983; Schidlowski, 1988; Mojzsis *et al.*, 1996; Rosing, 1999; Czaja *et al.*, 2013; Nutman *et al.*, 2016 |
| Origin of Methanogenesis | >2700 | Eigenbrode & Freeman, 2006 |
| Origin of Cyanobacteria | >2450 | Bekker *et al.*, 2004 |
| Origin of Eukaryotes | >1700 | Pang *et al.*, 2013 |
| Origin of plastids/Rhodophytes diverge | >1050 | Gibson *et al.*, 2018 |
| Akinetes diverge from cyanobacteria lacking cell differentiation | >1000 | Pang *et al.*, 2018 |

We created chronograms using both the uncorrelated gamma (UGAM) model (Drummond *et al*., 2006) and the autocorrelated CIR relaxed clock model (Lepage *et al*., 2007) to compare the effects of clock model type on the results. Two chains were run in parallel for each set of parameters, so that the two concurrent runs could be compared to one another as a test of convergence. Convergence of the MCMC chains was checked visually by plotting the summary statistics, and quantitatively by comparing the posterior distributions of two parallel chains using the *tracecomp* and *bpcomp* programs in PhyloBayes. We required an effective size >100 and a maximum difference between chains of <0.3. Simultaneous chains were run for approximately 36,000 cycles. Chronograms were generated using the *readdiv* function in Phylobayes 4.1, with approximately 20% of initial cycles discarded as burn-in. Chronograms were visualized using the phytools package in R (Revell, 2012). In order to test for the influence of the priors, we generated additional chronograms in the absence of sequence data using the - prior flag in Phylobayes. These chronograms displayed substantially different node timings, demonstrating that the priors did not overly influence the inferred dates and the sequence data informed the chronogram. Results from molecular clock analyses should be interpreted with caution, given the limitations associated with these analyses, including but not limited to changing generation times, the influence of natural selection, and variation in mutational rates across species (Ayala, 1999; Schwartz & Maresca, 2006; Bromham *et al*., 2018). Here, we attempted to ameliorate some of these challenges by using methods allowing for variation of rates between and across lineages, and by comparing results produced by different clock models. Finally, our goal with this analysis was not to pinpoint exact dates for many of the transitions discussed here, but rather to compare the relative timing on broad evolutionary scales.

### Identification of nitrogen-cycling genes and construction of gene trees

Curated gene database queries for each of the nitrogen-cycling genes we investigated were generated based on KEGG orthologies (Ogata *et al*., 1999) and downloaded from the UniProt database (The Uniprot Consortium, 2017). Nitrogen-cycling amino acid sequences for the gene trees were identified by conducting BLASTP (Altschul *et al*., 1990) searches of the open reading frames (ORFs) of every genome from the collection of 254 genomes. All ORFs were identified using Prodigal (Hyatt *et al*., 2010). The maximum e-value cutoff for BLAST hits was 10^−12^ and matches were excluded if the length of the local alignment was less than 50% of the length of the query sequence. BLAST results were compared with results from AnnoTree (Mendler *et al*., 2019) to verify gene distributions. Given that *nifK* and *nifD* have a shared evolutionary history (Fani *et al*., 2000), an e-value cutoff of 1e-30 was selected based on close examination of blastp hit results and KEGG annotations, which was sufficient to distinguish these subunits. For nitrogenase subunits *nifH, vnfD, vnfK, anfD, and anfK* we manually curated alignments by identifying key residues that were crucial for enzyme structure and function, as determined through literature searches and visualization using PyMol (Brigle *et al*., 1987; Kaiser *et al*., 2011; McGlynn *et al*., 2012; Howard *et al*., 2013; Keable *et al*., 2018). Jalview (Waterhouse *et al*., 2009) was used for visualization of key residues in alignments. All genes that did not include the key residues were removed from the alignment. Finally, to ensure that only the genes of interest were included in alignments and to verify annotations, all genes identified in the BLAST search were compared to the KEGG database using Kofam Koala (Aramaki *et al*., 2019), which assigns KEGG Ortholog numbers to each gene by using a homology search against a database of profile hidden Markov models. Only genes verified to be the gene of interest according to Kofam Koala were retained for downstream analysis. It is important to note that gene identification is necessarily limited by the search tools and databases used for annotation, and the methods used here were chosen to be conservative so as to ensure the removal of non-target genes from the analysis.

Alignments for the gene trees were created using MUSCLE (Edgar, 2004) and trimmed with TrimAl (Capella-Gutierrez *et al*., 2009) using the -automated1 option. The model of evolution was selected using Model Selection as implemented in IQTREE (Kalyaanamoorthy *et al*., 2017) using the default parameters. Trees were generated with RAxML-NG (Kozlov *et al*., 2019) using the model of evolution identified in IQ-TREE. Trees were run with at least 1000 bootstraps or until the diagnostic statistic based on the MRE-based bootstrapping test as implemented in RAxML-NG dropped below a cutoff of 0.03.

### Gene Tree and Species Chronogram Reconciliation

Gene trees were reconciled with species chronograms using the Analyzer of Gene and Species Trees (AnGST) (David & Alm, 2011). AnGST compares the topology of the gene tree with that of the species tree, rather than rely solely on presence and absence patterns, in order to identify gene birth, transfer, duplication, and loss events. Event penalties were set to hgt: 3, dup: 2, los: 1, and spc: 0. Ultrametric was set to True in order to constrain events temporally. Each run was conducted with 100 gene tree bootstraps in order to increase accuracy (David & Alm, 2011). Individual event timings were defined as the midpoint of the temporal region during which a given event could occur.

## Results

### Generation of a species tree and fossil-calibrated chronogram

We compared all gene trees to a species tree constructed from an alignment of concatenated sequences of 16 single-copy universal proteins from 308 organisms (Figure 2; genome list available as Supplemental Data). Monophyly was preserved for most major phyla, with three exceptions: (i) Tenericutes is contained within Firmicutes, (ii) the PVC superphylum contains Omnitrophica, and (iii) Lentisphaerae is nested within Verrucomicrobia. Additionally, in contrast to a recently published comprehensive tree of life (Hug *et al*., 2016), our tree does not place the recently-discovered Candidate Phyla Radiation (CPR) bacteria as the deepest-rooted bacterial clade; however, the CPR is placed as a sister group to the Cyanobacteria and Melainabacteria, which is consistent with that phylogeny (Hug *et al*., 2016). For the purposes of this study, placement of these groups should not greatly affect our results due to the high number of duplication/loss/transfer events we inferred overall across all groups. The species tree used for this analysis is a three-domain tree, in contrast to recent studies which have shown the addition of the Asgard Archaea to the tree of life to cause Eukaryotes to group within Archaea (Zaremba-Niedzwiedzka *et al*., 2017). However, we are agnostic as to the placement of the Asgard superphylum and the eukaryotes on the tree of life, as this was not our focus; the relationship of the three domains to one another should not substantially affect the results shown here due to the relative infrequency of inter-domain HGT relative to interdomain HGT, and the exclusion of eukaryotic nitrogen-cycling genes.

**Figure 2.**
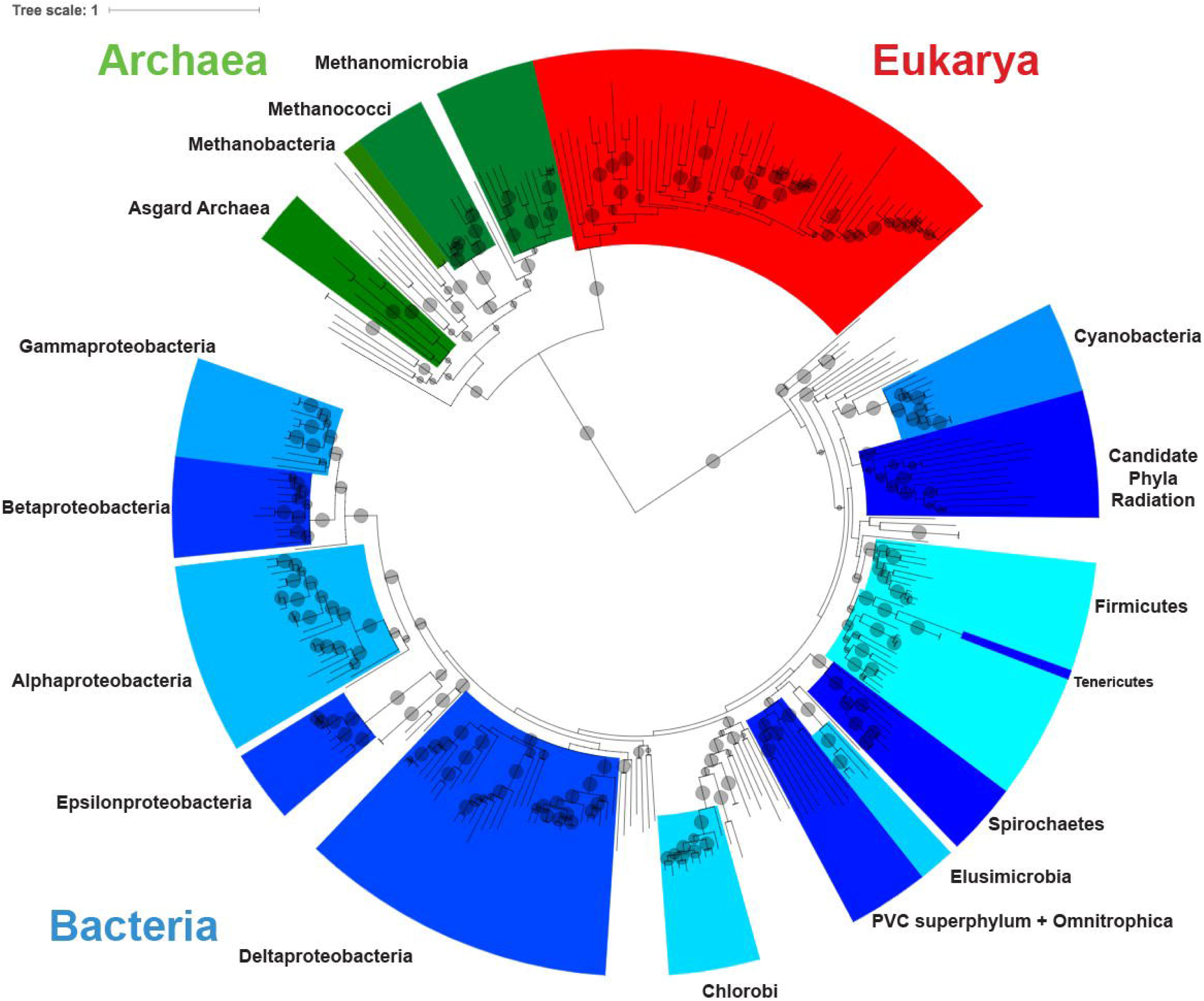
Species tree used for phylogenetic analysis. Maximum likelihood phylogeny based on an alignment of concatenated single-copy universal proteins from 308 genomes. Well-represented bacterial and archaeal phyla are labelled in black text. Bacterial clades are labeled in alternating colors of blue, archaea in green, eukaryotes in red. Bootstrap values (from 100 bootstraps) greater than 50 are shown as transparent gray circles, with larger circles representing higher bootstrap values.

We constructed four different chronograms from our species tree using two different clock models (UGAM and CIR) as well as liberal (representing the earliest date for which there is any evidence of a given event based on the current scientific literature) and conservative (representing the earliest date for which there is the most consensus for a given event based on the current scientific literature) fossil calibration points (see Methods). The liberal calibration points yielded an unreasonable root age (>4.5 Ga) and so were not further used for analysis. The UGAM clock model yielded a greater spread in estimated ages for node divergences, with an earlier root age (approximately 4044.68 +/- 143.093 Mya compared to 3982 +/- 131.218 Mya for the CIR clock model). The results shown in Figure 3 and Table 4 derive from the CIR clock model as this has been previously shown to outperform uncorrelated models (Lepage *et al*., 2007), but the results from the UGAM clock model are shown in Supplementary Figure 3 and Supplementary Table 2. All genome lists, alignments, and Newick files have been deposited in FigShare at: https://figshare.com/projects/Radiation_of_nitrogen_cycling_genes_across_the_tree_of_life/87461

**Table 2.** Number of nitrogen-cycling genes identified within 254 bacterial and archaeal genomes that were included in the analysis. All genes were identified from ORFs using blastp with an e-value cutoff of 1e-12, then filtered using the Kofam Koala tool. Nitrogenases were further refined by identifying key residues (see Methods).

| Gene | Number of genes identified |
| --- | --- |
| *amo* | 4 |
| *anfD* | 11 |
| *anfK* | 10 |
| *hao* | 0 |
| *napA* | 37 |
| *narG* | 34 |
| *nasA* | 28 |
| *nifD* | 165 |
| *nifK* | 92 |
| *nifH* | 233 |
| *nirB* | 39 |
| *nirK* | 10 |
| *nirS* | 16 |
| *norB* | 36 |
| *nosZ* | 21 |
| *nrfA* | 48 |
| *nxrA* | 34 |
| *vnfD* | 6 |
| *vnfK* | 6 |

**Table 3.** Frequencies of events inferred by AnGST for each gene as derived from both the CIR and UGAM clock models.

|  | CIR Clock Model |  |  |  | UGAM Clock Model |  |  |  |
| --- | --- | --- | --- | --- | --- | --- | --- | --- |
| Gene | HGT | Speciation | Loss | Duplication | HGT | Speciation | Loss | Duplication |
| *anfD* | 7 | 7 | 4 | 0 | 7 | 7 | 4 | 0 |
| *anfK* | 6 | 7 | 4 | 0 | 6 | 7 | 4 | 0 |
| *napA* | 13 | 46 | 23 | 0 | 12 | 50 | 26 | 0 |
| *narG* | 15 | 29 | 15 | 4 | 14 | 32 | 17 | 4 |
| *nasA* | 5 | 27 | 6 | 1 | 5 | 27 | 6 | 1 |
| *nifD* | 48 | 176 | 60 | 0 | 51 | 169 | 56 | 0 |
| *nifK* | 35 | 103 | 48 | 1 | 34 | 107 | 51 | 1 |
| *nifH* | 68 | 224 | 72 | 12 | 74 | 202 | 56 | 12 |
| *nirB* | 14 | 34 | 13 | 3 | 13 | 38 | 16 | 3 |
| *nirK* | 4 | 9 | 5 | 1 | 4 | 9 | 5 | 1 |
| *nirS* | 7 | 13 | 6 | 1 | 7 | 13 | 6 | 1 |
| *norB* | 26 | 13 | 5 | 2 | 25 | 13 | 5 | 2 |
| *nosZ* | 13 | 5 | 0 | 2 | 13 | 5 | 0 | 2 |
| *nrfA* | 24 | 34 | 13 | 2 | 23 | 37 | 15 | 2 |
| *nxrA* | 13 | 34 | 18 | 4 | 13 | 34 | 18 | 4 |
| *vnfD* | 3 | 3 | 1 | 0 | 3 | 3 | 1 | 0 |
| *vnfK* | 3 | 3 | 1 | 0 | 3 | 3 | 1 | 0 |

**Table 4.** Inferred birth dates for nitrogen-cycling genes based on the chronogram generated using the CIR clock model with conservative calibration points. All gene birth events are inferred to occur between nodes on the species chronogram, and therefore our methods do not allow us to infer specific dates for gene birth events. Here we list the inferred timing for the earliest possible timing (upper node) and latest possible timing (lower node) with 95% confidence intervals as reported by PhyloBayes, as well as the calculated midpoint between the two node times. Times are reported in millions of years ago. The geological era reported in the right column corresponds to the midpoint between the upper and lower node. Given the uncertainty inherent in the estimation of node timings and inference in events, these are provided for geological reference and should not be taken as definitive timings. The equivalent table, generated using the UGAM clock model, is depicted as Supplementary Table 2.

| Gene | Upper node date | 95% confidence interval | Lower node date | 95% confidence interval | Midpoint between nodes | Geologic era |
| --- | --- | --- | --- | --- | --- | --- |
| *vnfD* | 1445.24 | 1052.37, 1735.83 | 482.23 | 210.11, 778.444 | **963.73** | Neoproterozoic |
| *vnfK* | 1445.24 | 1052.37, 1735.83 | 482.23 | 210.11, 778.444 | **963.73** | Neoproterozoic |
| *nosZ* | 1325.57 | 1048.93, 1588.52 | 1139.65 | 851.174, 1408.81 | **1232.61** | Mesoproterozoic |
| *nirK* | 1452.66 | 1193.07, 1706.53 | 1343.08 | 1094.21, 1585.92 | **1397.87** | Mesoproterozoic |
| *nirS* | 1913.87 | 1694.13, 2144.31 | 1423.17 | 1153.32, 1677.23 | **1668.52** | Paleoproterozoic |
| *norB* | 1913.87 | 1694.13, 2144.31 | 1423.17 | 1153.32, 1677.23 | **1668.52** | Paleoproterozoic |
| *anfD* | 2349.58 | 2153.39, 2583.93 | 1745.53 | 1498.39, 1999.54 | **2047.55** | Paleoproterozoic |
| *anfK* | 2349.58 | 2153.39, 2583.93 | 1745.53 | 1498.39, 1999.54 | **2047.55** | Paleoproterozoic |
| **nasA** | 2540.50 | 2351.4,<br>2778.03 | 2063.23 | 1848.31,<br>2293.91 | **2301.87** | **Paleoproterozoic** |
| **nrfA** | 2454.68 | 2178.51,<br>2737.37 | 2267.54 | 1918.96,<br>2574.71 | **2361.11** | **Paleoproterozoic** |
| **nirB** | 2748.37 | 2545.88,<br>2989.62 | 2603.25 | 2384.99,<br>2850.17 | **2675.81** | **Neoarchean** |
| **nifK** | 2926.54 | 2734.92,<br>3180.11 | 2629.22 | 2415.05,<br>2877.89 | **2777.88** | **Neoarchean** |
| **nxrA** | 2816.89 | 2630.27,<br>3063.26 | 2787.46 | 2603.09,<br>3029.43 | **2802.17** | **Mesoarchean** |
| **napA** | 2816.89 | 2630.27,<br>3063.26 | 2787.46 | 2603.09,<br>3029.43 | **2802.17** | **Mesoarchean** |
| **narG** | 2816.89 | 2630.27,<br>3063.26 | 2787.46 | 2603.09,<br>3029.43 | **2802.17** | **Mesoarchean** |
| **nifD** | 2934.72 | 2744.91,<br>3187.67 | 2869.82 | 2682.33,<br>3118.74 | **2902.27** | **Mesoarchean** |
| **nifH** | 3060.98 | 2863.64,<br>3325.02 | 3026.38 | 2832.29,<br>3286.79 | **3043.68** | **Mesoarchean** |

**Figure 3.**
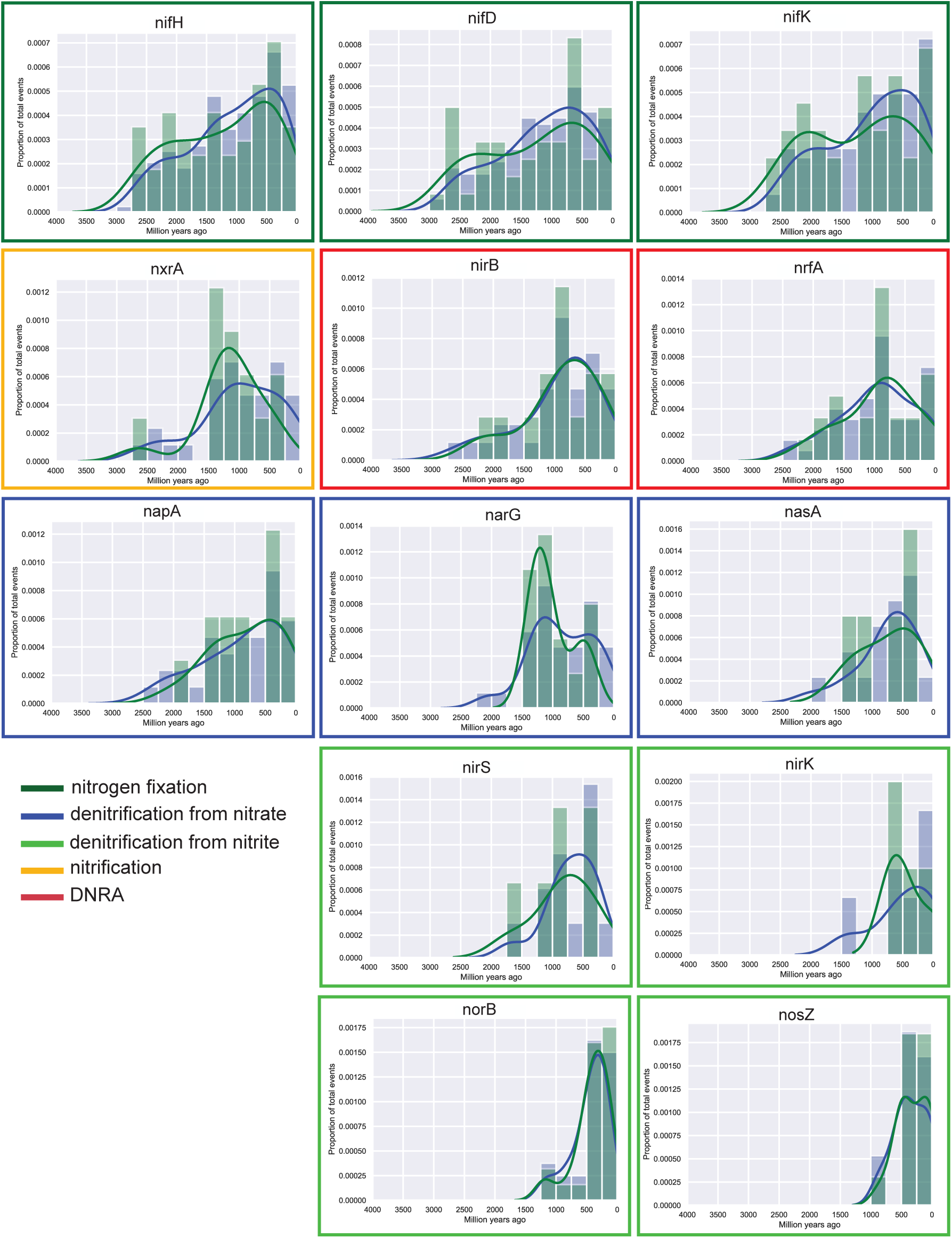
Histograms and density plots of all speciation, loss, duplication and horizontal gene transfer (HGT) events. All events are shown in blue, HGT events alone are shown in green. Events were inferred by reconciling phylogenetic trees of nitrogen-cycling genes with a chronogram generated using a CIR clock model and conservative calibration points. Y-axis represents proportion of all events (blue) or HGT events only (green). Borders around graphs indicate the metabolic process that the gene is involved in. Histograms are ordered according to the earliest time point at which a speciation, loss, duplication, or HGT event occurred. An equivalent figure with dates inferred from the UGAM clock model is shown in Supplementary Figure 3.

### Identification of duplication, speciation, loss, and horizontal gene transfer events for nitrogen-cycling genes

We identified 18 different nitrogen-cycling genes for analysis of horizontal gene transfer, speciation, duplication, and loss events (Table 2). This methodology is best suited to genes that have many representatives from a diverse suite of taxa. Very few genes from the *hao* and *amo* gene families were identified in these genomes or passed our stringent filtration tests, and thus they did not yield sufficient data for robust conclusions and were not included in further analyses. We generated maximum likelihood gene trees for each of the remaining genes (Supplementary Data) and compared these gene trees to the fossil-calibrated chronogram in order to identify and infer the timing of birth, duplication, loss, and horizontal gene transfer events for each of these genes. The number of speciation events inferred for each gene in this study was relatively high compared to the number of inferred loss and HGT events (Table 3), with very few duplication events observed. Inferred loss events generally skewed younger than inferred horizontal gene transfer and speciation events. Our results indicate that the iron-molybdenum nitrogenases (*nifH/nifD/nifK*) were among the oldest genes, originating during the Archean era and beginning to spread across the tree of life via horizontal gene transfer fairly early in Earth history (Fig. 3). In contrast, the alternative nitrogenases *anf* and *vnf* were inferred to have arisen and radiated across the tree of life much later (Table 4, Supplementary Table 2, Supplementary Figures 4 and 5), but very few *anf* and *vnf* genes were identified among our sample set and therefore these results should be treated with caution (Table 2). The denitrifying genes *norB, nosZ, nirK* and *nirS* were inferred to have arisen later in Earth history (Table 4) and began to proliferate across the tree of life much more recently, up to approximately 1.5 Ga (Figure 3). These genes encode enzymes that catalyze denitrification processes from nitrite to nitric oxide, nitrous oxide, and dinitrogen gas. The genes *nrfA* and *nirB*, which are involved in the DNRA process, are inferred to have arisen by approximately 2.7-2.2 Ga, and began to increasingly proliferate widely across the tree of life into the Mesoproterozoic. Genes involved in nitrate reduction, including *narG, nasA*, and *napA*, were inferred to have arisen relatively early (approximately 2.8 Ga for *narG* and *napA*; approximately 2.3 Ga for *nasA*) (Table 4) and we inferred a few speciation and horizontal gene transfer events for these genes between 2-2.5 Ga, but we did not observe a rise in HGT events for these genes until much later, at approximately 1.5 Ga (Figure 3). We observed a similar trend for *nxrA*, a gene involved in nitrite oxidation, which was inferred to have arisen around 2.8 Ga but did not exhibit a rise in speciation, duplication, or transfer events until approximately 1.5 Ga (Figure 3, Table 4).

Taken together, we observed that genes related to the fixation of nitrogen from dinitrogen gas to ammonium arose early and proliferated across the tree of life relatively quickly, while genes related to nitrate reduction and nitrite oxidation also arose early but did not begin to proliferate across the tree of life until later. Genes related to denitrification, particularly downstream from nitrite, arose much later (Figure 3). Our results regarding the timing of the birth, duplication, speciation, loss and HGT events for specific nitrogen-cycling genes showed a few differences between the CIR (Figure 3, Table 4) and UGAM clock models (Supplementary Figure 3, Supplementary Table 2). However, the overall patterns in the relative timing for the birth and spread of specific genes in the nitrogen cycle were similar regardless of the type of clock models used.

## Discussion

We have focused here on tracking gene birth, duplication, speciation, and particularly horizontal gene transfer events across deep time. The acquisition of new functional genes is particularly important because it can allow clades of microbes to invade new ecological niches. The rate of horizontal gene transfer itself is likely to be related to variables like cell density, co-localization of donor and recipient, cell diversity, and the types of organisms involved (Gogarten & Townsend, 2005). Although some have argued for a neutral theory of gene transfer in which horizontally acquired genes are not adaptive (Gogarten & Townsend, 2005; Andreani *et al*., 2017), studies indicating that horizontally transferred genes perform crucial cellular functions and enable adaptation to specific ecological niches suggest that horizontally acquired genes are generally adaptive (Daubin & Ochman, 2004; Coleman & Chisholm, 2010; Burke *et al*., 2011; Popa *et al*., 2011; Polz *et al*., 2013; McInerney *et al*., 2017; Moulana *et al*., 2020). Moreover, genes providing selective advantages are more likely to be retained in the genome than genes acquired due to neutral transfer, which are more likely to be purged, thus strengthening the signal of adaptive HGT events in the genomic record. Therefore, any observations of a rise in the relative number of successful horizontal gene transfers for a given gene are likely to give an indication of the relative availability or metabolic importance of a given substrate for that gene. As such, studying the history of HGT of specific genes can provide insights into the points at which possession of such genes provided substantial selective advantages, which can then be used as a metric for when a specific metabolism became feasible or energetically favorable, or when the substrates of specific enzymes became relatively abundant. A crucial caveat to this method, however, is that it cannot infer relative abundance or population sizes of the organisms carrying these genes. Thus, if a specific strain carrying a nitrogen metabolizing gene grows in abundance with no transfer of genes to other lineages, our method would observe low rates of transfer for genes involved in that metabolism. Additionally, a poor phylogeny caused by weak phylogenetic signal within the alignment would make it difficult to precisely identify such horizontal gene transfer events. Therefore, these results, while easily overinterpreted, should largely be interpreted in the context of other studies, especially those which use substantially different methodologies.

On the whole, our results provide support for geochemical data indicating that biological nitrogen fixation was an important source of fixed nitrogen in the Archaean (Stüeken *et al*., 2015; Ossa Ossa *et al*., 2019). These results also suggest that local sources of nitrate may have been exploited by denitrifying microbes, while denitrifiers using nitrite or downstream products would not have proliferated until much later in Earth history, until the mid-Proterozoic, well after the Great Oxidation Event. This finding is consistent with previous geochemical studies that documented isotopic evidence of denitrification in the Neoarchean (Garvin *et al*., 2009b; Godfrey & Falkowski, 2009; Koehler *et al*., 2019) and Paleoproterozoic (Zerkle *et al*., 2017a; Kipp *et al*., 2018; Luo *et al*., 2018). Our results suggest that genes involved in nitrate reduction arose by approximately 2.8 Ga, indicating that nitrate was present and used as a metabolic substrate at that time. However, our data suggest that denitrification may not have been a reliable energy source until about 1.5 Ga, and therefore used by a smaller diversity of clades. From the mid-Proterozoic onwards, nitrate may have been sufficiently bioavailable to make it a more widely-used substrate. In the following, we discuss the implications of our results for the evolution of the nitrogen cycle and its relationship to the redox state of the Earth.

### Nitrogen Fixation

Our data indicate that nitrogen fixation through the use of molybdenum nitrogenase (Nif) is an ancient process, arising by approximately 3.1-2.7 Ga (Table 4). Our results are consistent with previous work from geochemical analyses suggesting that nitrogen fixation must have arisen early in order to support an expanding biosphere (Stüeken *et al*., 2016a; Koehler *et al*., 2019; Ossa Ossa *et al*., 2019). In contrast, a previous analysis based on the evolutionary rate of nitrogenase genes suggested that functional Mo-nitrogenase arose relatively late (approx. 2.2-1.5 Ga) (Boyd *et al*., 2011a), which may also be consistent with the hypothesis that modern planktonic nitrogen fixers did not become abundant in global oceans until the Neoproterozoic (Sánchez-Baracaldo *et al*., 2014). Our results, which are based on reconciliation of gene trees with a chronogram inferred from universally conserved, single-copy genes, are instead consistent with recent nitrogen isotope evidence for biological nitrogen fixation back to at least 3.2 Ga (Stüeken *et al*., 2015). It is also consistent with phylogenetics work suggesting that molybdenum nitrogenase arose early in the evolution of life on Earth (Raymond *et al*., 2004). Similarly, phylogenetic reconstructions showing that Mo-nitrogenases arose before V- and Fe-nitrogenases are consistent with this conclusion (Garcia *et al*., 2020). It has been argued that nitrogenase was present in LUCA (Weiss *et al*., 2016), though that has been disputed (Boyd *et al*., 2011a; Mus *et al*., 2019; Berkemer & McGlynn, 2020) and our results do not resolve this issue. However, the identification of horizontal gene transfer events for nitrogenase subunits during the Archean (Figure 3) suggest that this metabolism may have been abundant and beneficial enough during this time period to have been successfully transferred and retained in microbial genomes.

Our results support the argument that abiotic sources of fixed nitrogen were unlikely to be significant enough to sustain the early biosphere (Raymond *et al*., 2004; Canfield *et al*., 2010). It has been proposed that early life received its nitrogen from the lightning-catalyzed reaction between N_2_ and CO_2_ as a source of NO_x_, and that a steady reduction in atmospheric CO_2_ levels reduced this flux, eventually leading to a nitrogen crisis around 2.2 Ga that would have favored the rise and spread of biological nitrogen fixation (Navarro-González *et al*., 2001). In contrast, several studies have concluded that abiotic nitrogen fixation on the early Earth would produce 50-to 5000-fold lower rates of fixed nitrogen than what is contained in the modern ocean, making nitrogen highly limiting to the early biosphere, even at higher CO_2_ levels (Canfield *et al*., 2010). Our data suggest that biological nitrogen fixation arose and proliferated early and that acquisition of the nitrogenase enzyme provided enough of a selective advantage to be successfully transferred across lineages during the early Archaean, supporting the notion that fixed nitrogen was not widely available.

Moreover, if the radiation of biological N_2_ uptake dates back to the early Archean, it may have had a significant impact on atmospheric pressure. Today, nitrogen makes up approximately 78% of the Earth’s modern atmosphere by volume, predominantly as N_2_ gas. The atmosphere of the early Earth was probably also N_2_-rich, but while some models suggest a 2-4 times larger atmospheric N_2_ reservoir in the Archean (Johnson & Goldblatt, 2018), proxy evidence ranges from near modern values (Marty *et al*., 2013; Avice *et al*., 2018) to less than half of today’s reservoir (Som *et al*., 2016). If atmospheric N_2_ pressure changed over time, it is likely that biological activity would have played a major role in driving these changes by enhancing nitrogen burial in sediments and by accelerating oxidative weathering of nitrogen from continental crust (Stüeken *et al*., 2016b; Zerkle & Mikhail, 2017). If nitrogen was fixed by nitrogenase early in the Archaean, as indicated by our results, then biological nitrogen burial began long before the onset of oxidative weathering in the Neoarchean (Stüeken *et al*., 2012) which makes it possible that atmospheric N_2_ decreased over the course of the Archean. Although further paleobarometric proxies are needed to verify this trend, our results provide an important anchor point for the onset of biological N_2_ drawdown.

Lastly, though we identified very few iron and vanadium nitrogenases (*anf* and *vnf*, respectively) in our datasets, our results tentatively suggest that these nitrogenases radiated across the tree of life later than the iron-molybdenum nitrogenases (*nif*), which supports conclusions from other phylogenetics studies (Garcia *et al*., 2020). The early rise of Mo-dependent nitrogenase would have required a source of Mo to act as a cofactor for Mo-containing nitrogenases, implying that nanomolar levels of dissolved Mo (Scott *et al*., 2008; Reinhard *et al*., 2013), presumably derived from anoxic weathering or hydrothermal sources of molybdenum to the early ocean, were sufficient for the development of Nif and the proliferation of nitrogen fixers. Oxidative weathering was therefore evidently not required for the onset of biological nitrogen fixation (c.f. Boyd *et al*., 2011b). We speculate that the evolutionary pressure for the diversification of alternative nitrogenases arose in the Neoproterozoic with the rise of eukaryotic algae (Brocks *et al*., 2017), which may have substantially increased the nitrogen demand in the global ocean. Vanadium and iron-based nitrogenases are less efficient at fixing nitrogen than Nif (Mus *et al*., 2018), which perhaps lowers the probability of successful gene transfer for Vnf and Anf. However, when nitrogen-demand increased in the environment, it is conceivable that more of these genes were shared successfully. Another hypothesis to explain our data is that the gene transfer of alternative nitrogenases was affected by global climate. Vnf becomes more efficient than Nif at cold temperatures (Miller & Eady, 1988) and the activity of Nif-using cyanobacteria decreases at high latitudes (Brauer *et al*., 2013). Thus, the cold climate of the Cryogenian’Snowball Earth’ period (720-635 Ma) (Hoffman *et al*., 2017) may have increased the fitness of organisms possessing Vnf. However, these hypotheses remain speculative until more genomic data for Vnf and Anf has been acquired. In any case, it is important to keep in mind that the origin of alternative nitrogenases likely occurred long before the Neoproterozoic. Vanadium nitrogenase in *Azotobacter vinelandii* has been shown to reduce carbon monoxide (CO) to the hydrcarbons ethylene (C_2_H_2_), ethane (C_2_H_6_), and propane (C_3_H_8_) (Lee *et al*., 2010), which supports the origin of V-nitrogenase in the Archean eon when CO was more abundant in the atmosphere (Anbar & Knoll, 2002). Additionally, V was bioavailable in Archean ocean waters under slightly acidic conditions (Moore *et al*., 2020). The genomes of more V-nitrogenase and Fe-nitrogenase organisms must be sequenced to better understand the evolutionary history of these alternative nitrogenases.

### Nitrogen Cycling Prior to the Oxygenation of the Oceans and Atmosphere

Although our results suggest that nitrogen fixation was an important process on the early Earth, microbial metabolisms making use of more oxidized forms of nitrogen, particularly through the process of denitrification downstream from nitrite, do not appear to have arisen and spread across the tree of life until much later. Some early steps in the nitrogen cycle, such as nitrate reduction to nitrite via *nasA, narG* or *napA*, appear to have arisen relatively early (in the late Archean or early Proterozoic), consistent with geochemical record of denitrification at that time (Godfrey & Falkowski, 2009; Koehler *et al*., 2019). However, we did not identify many horizontal gene transfer events early in the evolutionary history of these genes, and their proliferation across the tree of life only began in earnest much later, approximately 1.5 Ga. Our results therefore suggest that nitrate availability was restricted in space and/or time. Indeed, thermodynamic constraints and environmentally resolved geochemical datasets show that the anoxic deep ocean of the Precambrian contained ammonium while nitrate was restricted to surface waters (Stüeken *et al*., 2016a; Yang *et al*., 2019). It is therefore likely that the nitrate-reducing genes appeared in locally oxygenated regions of the Archean surface ocean, which have been inferred from isotopic studies back to 3.0 Ga (Olson *et al*., 2013; Planavsky *et al*., 2014). In such oxygen oases, oxygen concentrations may have reached micromolar concentrations (Olson *et al*., 2013), which is high enough for the complete oxidation of ammonium to nitrite and nitrate (Lipschultz *et al*., 1990). Alternatively, it a small flux of nitrogen oxides from lightning and volcanism may have existed (Navarro-González *et al*., 2001; Mather *et al*., 2004) and maintained populations of nitrate-reducing denitrifiers. Denitrification is a highly energy-yielding metabolism (Schoepp-Cothenet *et al*., 2013), and therefore even a small source flux of nitrate to the early ocean would probably have been exploited. However, we stress that denitrification left its geochemical mark in only a few Archean basins (Garvin *et al*., 2009b; Godfrey & Falkowski, 2009; Koehler *et al*., 2019), while other localities show no geochemical evidence of nitrate reduction (Ossa Ossa *et al*., 2019). This observation may argue against a significant contribution of lightning-derived nitrogen oxides, which we would expect to have been more uniformly distributed.

Our results suggest that genes for nitrite-metabolizing enzymes were not frequently transferred during the early Archean, which may imply that nitrite-metabolizing genes were not widespread in Archean ecosystems. It is possible that only specific taxa were responsible for the conversion of nitrite into nitrate or ammonium. Alternatively, the conversion of nitrite to ammonium may have happened abiotically. Ferrous iron (Fe^2+^), which is thought to have been abundant in the early anoxic ocean, may have readily reduced nitrite to ammonium or nitrogen gas without biological intervention (Summers & Chang, 1993; Canfield *et al*., 2010). Nitrite levels during this time period were therefore likely too low to cause substantial proliferation of genes for enzymes which take nitrite as a substrate, while nitrate and ammonium concentration were high enough for selection to favor biological metabolism of these molecules by diverse microorganisms.

### Effects of Increased Oxygen Levels on the Nitrogen Cycle

Multiple lines of evidence point to increasing oxygenation of surface environments from 2.75 Ga onwards, culminating in the Great Oxidation Event (GOE) at 2.3 Ga (Noffke *et al*., 2007; Crowe *et al*., 2013; Lyons *et al*., 2014; Planavsky *et al*., 2014). Our data suggest that the genes *nrfA* and *nirB*, which reduce a variety of oxidized forms of nitrogen to ammonium through the DNRA pathway, arose in the Neoarchean or Paleoproterozoic, but the relative number of HGT events for this gene began to rise approximately around the time of the GOE. Unlike denitrification to N_2_ gas, DNRA retains the reduced nitrogen product in the system as aqueous NH_4_^+^, which may have prevented a potential ‘nitrogen crisis’ during the wake of the GOE (Falkowski & Godfrey, 2008). The isotopic fingerprint of DNRA can so far not be distinguished from denitrification in the geochemical record, and thus our results provide the first indication that this metabolism may have been of greater ecological importance than previously proposed. DNRA rather than denitrification may therefore explain geochemical evidence of biological nitrate reduction in the Neoarchean and early Proterozoic (Garvin *et al*., 2009b; Godfrey & Falkowski, 2009; Zerkle *et al*., 2017b; Kipp *et al*., 2018), i.e. long before our observed radiation in denitrification genes. This observation may suggest that these early nitrate-reducing ecosystems were primarily performing DNRA, consistent with iron-rich conditions in the Archean and Paleoproterozoic ocean (Michiels *et al*., 2017).

After approximately 1.5 Ga, the relative number of HGT events for enzymes involved in the modern denitrification pathway began to rise. Additionally, our data suggest that the gene *nxr*, which oxidizes nitrite into nitrate, arose relatively early (approx. 2.8 Ga) but did not begin to spread across the tree of life until ∼1.5 Ga. One possibility is that increased copper availability facilitated the spread of nitrification/denitrification, which require enzymes that rely on copper cofactors (Moore *et al*., 2017). However, geochemical evidence of increasing copper concentrations in seawater at 1.5 Ga is so far lacking. Another possibility is that the expansion of these metabolisms is linked to a Mesoproterozoic rise in oxygen, which has recently been proposed in several geochemical studies (Cox *et al*., 2016; Canfield *et al*., 2018; Zhang *et al*., 2018; Shang *et al*., 2019) Higher oxygen levels would have led to an expansion of the marine nitrate reservoir, which may in turn have been critical for the rise of eukaryotes around this time (Anbar & Knoll, 2002; Knoll & Nowak, 2017).

The primary substrate for many of the enzymes that demonstrated this late rise in the number of HGT events is nitrite, suggesting that nitrite levels may have increased at this time, such that selection favored the spread of nitrite-metabolizing enzymes across the tree of life. Nitrite is thermodynamically unstable in oxic conditions, but it occurs transiently in chemoclines in the modern ocean because it is abundantly produced as an intermediate in nitrification and denitrification pathways (Wada & Hatton, 1971). A more vigorous aerobic nitrogen cycle in the Proterozoic may thus be linked to the establishment of a more permanent dynamic nitrite reservoir. Abiotic reduction of nitrite by Fe^2+^, which was perhaps dominant during the Archean (see above), may have slowed down with the deepening of the chemocline after the GOE.

Several caveats must be kept in mind while interpreting these results. As mentioned above, the accuracy of molecular clocks is constrained by the models used to infer dates and the accuracy of the fossil-based time points used for clock calibration. We sought to minimize the influence of these parameters by using both liberal and conservative time points and by employing two different clock models, but nevertheless these limitations must be taken into consideration, and thus we emphasize the relative rather than the absolute timing of the events identified here. Importantly, the results generated through the use of liberal calibration points are not intended to be taken as an older bound for these gene proliferations, but rather as a test of the sensitivity of our results to the specifications of the molecular clock. Moreover, the methods used to identify genes used in the analysis were designed to be conservative so as to reduce the possibility that spurious genes were included in the analysis, but these methods are inherently limited by the quality of the databases used for annotation. These conservative methods precluded the inclusion of *hao* and *amo*, which are involved in the oxidation of ammonia to nitrite and nitrogen gas, in our analysis. Others have observed that archaeal ammonia oxidizers most likely originated during the GOE and began to spread through the shallow ocean approximately 800 Mya (Ren *et al*., 2019). As new genes and genomes continue to be sequenced and analyzed for gene function, the quality of such analyses will improve over time.

## Conclusion

Our results show that biological nitrogen fixation appears to have arisen and proliferated early in the Archaean, and that genes in the denitrification pathway and genes related to the consumption of nitrite and its downstream products began to proliferate across the tree of life following the oxygenation of Earth’s atmosphere and oceans. Moreover, our results support the hypothesis that the molybdenum-based variety of nitrogenase diversified much earlier than alternative nitrogenases, which may only have become important with the increasing nitrogen demand of algae in the Neoproterozoic, implying that molybdenum was not a limiting resource in the Archean ocean. Furthermore, our data provide the first indirect evidence for a small source of nitrate to the early ocean, possibly as a result of lightning and volcanism or from localized oxygen oases. Some of this nitrate appears to have been used for nitrate reduction to ammonium (DNRA), which may have helped overcome nitrogen limitation by retaining fixed nitrogen in the system. DNRA rather than denitrification may explain some of the isotopic records of biological nitrate utilization in the Archean and early Proterozoic. We cannot confirm the hypothesized suppression of N_2_O-metabolizing enzymes in the mid-Proterozoic, but our results are consistent with vigorous nitrification and denitrification after the GOE, in particular from 1.5 Ga onwards, which may have led to leakage of N_2_O into the atmosphere and the stabilization of global climate. The proliferation of key nitrogen-metabolizing genes across the tree of life at different points in Earth’s redox history provides important insights into the evolution and radiation of microbial metabolisms on the early Earth in response to major environmental transitions and supports the notion that increasing nitrate levels in the mid-Proterozoic may have contributed to the rise of eukaryotic life.

## Acknowledgements

We thank three anonymous reviewers for extremely detailed and helpful comments that greatly improved the manuscript. We would like to thank Rou-Jia Sung for assistance with structural predictions, as well as Casey Bryce, Colin Goldblatt, Ben Johnson, John Baross, Victoria Meadows, Jaclyn Saunders, and Cara Magnabosco for support and useful discussions. This work was performed as part of NASA’s Virtual Planetary Laboratory, supported by the NASA Astrobiology Program under grant 80NSSC18K0829 as part of the Nexus for Exoplanet System Science (NExSS) research coordination network. CWP received a Towsley Fellowship from Carleton College, CR was supported by the Virtual Planetary Laboratory, KM was supported by a Student Research Partnership grant from Carleton College, and REA was supported in part by a NASA Postdoctoral Fellowship from the NASA Astrobiology Institute.

## Notes

### Competing Interest Statement

The authors have declared no competing interest.

### Summary of Updates

Tables 3 and 4 have been updated, and the manuscript has been edited to reflect this.

## References

Altschul SF, Gish W, Miller W, Myers EW, Lipman DJ (1990) Basic local alignment search tool. Journal of Molecular Biology 215, 403–410.

Anbar AD, Knoll AH (2002) Proterozoic ocean chemistry and evolution: A bioinorganic bridge? Science 297, 1137–1142.

Andreani NA, Hesse E, Vos M (2017) Prokaryote genome fluidity is dependent on effective population size. The ISME Journal 11, 1719–1721.

Aramaki T, Blanc-Mathieu R, Endo H, Ohkubo K, Kanehisa M, Goto S, Ogata H (2019) KofamKOALA: KEGG ortholog assignment based on profile HMM and adaptive score threshold. Bioinformatics.

Avice G, Marty B, Burgess R, Hofmann A, Philippot P, Zahnle K, Zakharov D (2018) Evolution of atmospheric xenon and other noble gases inferred from Archean to Paleoproterozoic rocks. Geochimica et Cosmochimica Acta 232, 82–100.

Ayala FJ (1999) Molecular clock mirages. BioEssays 21, 71–75.

Beiko RG, Harlow TJ, Ragan MA (2005) Highways of gene sharing in prokaryotes. Proceedings of the National Academy of Sciences of the United States of America 102, 14332–7.

Bekker A, Holland HD, Wang P-L, Rumble D, Stein HJ, Hannah JL, Coetzee LL, Beukes NJ (2004) Dating the rise of atmospheric oxygen. Nature 427, 117–120.

Berkemer SJ, McGlynn SE (2020) A New Analysis of Archaea–Bacteria Domain Separation: Variable Phylogenetic Distance and the Tempo of Early Evolution. Molecular Biology and Evolution.

Boyd ES, Anbar AD, Miller S, Hamilton TL, Lavin M, Peters JW (2011a) A late methanogen origin for molybdenum-dependent nitrogenase. Geobiology 9, 221–32.

Boyd ES, Hamilton TL, Peters JW (2011b) An alternative path for the evolution of biological nitrogen fixation. Frontiers in Microbiology 2, 1–11.

Boyd ES, Peters JW (2013) New insights into the evolutionary history of biological nitrogen fixation. Frontiers in microbiology 4, 201.

Brandes JA, Boctor NZ, Cody GD, Cooper BA, Hazen RM, Yoder HS (1998) Abiotic nitrogen reduction on the early Earth. Nature 395, 365–367.

Brauer VS, Stomp M, Rosso C, Beusekom SA van, Emmerich B, Stal LJ, Huisman J (2013) Low temperature delays timing and enhances the cost of nitrogen fixation in the unicellular cyanobacterium Cyanothece. The ISME Journal 7, 2105–2115.

Brigle KE, Weiss MC, Newton WE, Dean DR (1987) Products of the iron-molybdenum cofactor-specific biosynthetic genes, nifE and nifN, are structurally homologous to the products of the nitrogenase molybdenum-iron protein genes, nifD and nifK. Journal of bacteriology 169, 1547–53.

Brocks JJ, Jarrett AJM, Sirantoine E, Hallmann C, Hoshino Y, Liyanage T (2017) The rise of algae in Cryogenian oceans and the emergence of animals. Nature 548, 578–581.

Bromham L, Duchêne S, Hua X, Ritchie AM, Duchêne DA, Ho SYW (2018) Bayesian molecular dating: Opening up the black box. Biological Reviews.

Buick R (2007) Did the Proterozoic “Canfield Ocean” cause a laughing gas greenhouse? Geobiology 5, 97–100.

Burke C, Steinberg P, Rusch D, Kjelleberg S, Thomas T (2011) Bacterial community assembly based on functional genes rather than species. Proceedings of the National Academy of Sciences of the United States of America 108, 14288–14293.

Canfield D, Kristensen E, Thamdrup B (2005) Aquatic Geomicrobiology, Volume 48.

Canfield DE, Glazer AN, Falkowski PG (2010) The evolution and future of Earth’s nitrogen cycle. Science 330, 192–6.

Canfield DE, Zhang S, Frank AB, Wang X, Wang H, Su J, Ye Y, Frei R (2018) Highly fractionated chromium isotopes in Mesoproterozoic-aged shales and atmospheric oxygen. Nature Communications 9, 2871.

Capella-Gutierrez S, Silla-Martinez JM, Gabaldon T (2009) trimAl: a tool for automated alignment trimming in large-scale phylogenetic analyses. Bioinformatics 25, 1972–1973.

Coleman ML, Chisholm SW (2010) Ecosystem-specific selection pressures revealed through comparative population genomics. Proceedings of the National Academy of Sciences of the United States of America 107, 18634–9.

Cox GM, Jarrett A, Edwards D, Crockford PW, Halverson GP, Collins AS, Poirier A, Li Z-X (2016) Basin redox and primary productivity within the Mesoproterozoic Roper Seaway. Chemical Geology 440, 101–114.

Crowe SA, Døssing LN, Beukes NJ, Bau M, Kruger SJ, Frei R, Canfield DE (2013) Atmospheric oxygenation three billion years ago. Nature 501, 535–538.

Czaja AD, Johnson CM, Beard BL, Roden EE, Li W, Moorbath S (2013) Biological Fe oxidation controlled deposition of banded iron formation in the ca. 3770 Ma Isua Supracrustal Belt (West Greenland). Earth and Planetary Science Letters 363, 192–203.

Darling AE, Jospin G, Lowe E, Matsen FA, Bik HM, Eisen JA (2014) PhyloSift: phylogenetic analysis of genomes and metagenomes. PeerJ 2, e243.

Daubin V, Ochman H (2004) Bacterial Genomes as New Gene Homes: The Genealogy of ORFans in E. coli. Genome Research 14, 1036–1042.

David LA, Alm EJ (2011) Rapid evolutionary innovation during an Archaean genetic expansion. Nature 469, 93–6.

Delmont TO, Quince C, Shaiber A, Esen ÖC, Lee ST, Rappé MS, MacLellan SL, Lücker S, Eren AM (2018) Nitrogen-fixing populations of Planctomycetes and Proteobacteria are abundant in surface ocean metagenomes. Nature Microbiology 3, 804–813.

Drummond AJ, Ho SYW, Phillips MJ, Rambaut A (2006) Relaxed Phylogenetics and Dating with Confidence. PLoS Biology 4, e88.

Edgar RC (2004) MUSCLE: multiple sequence alignment with high accuracy and high throughput. Nucleic Acids Research 32, 1792–1797.

Eigenbrode JL, Freeman KH (2006) Late Archean rise of aerobic microbial ecosystems. Proceedings of the National Academy of Sciences of the United States of America 103, 15759–15764.

Falkowski PG, Godfrey L V (2008) Electrons, life and the evolution of Earth’s oxygen cycle. Philosophical Transactions of the Royal Society B: Biological Sciences 363, 2705–2716.

Fani R, Gallo R, Liò P (2000) Molecular Evolution of Nitrogen Fixation: The Evolutionary History of the nifD, nifK, nifE, and nifN Genes. Journal of Molecular Evolution 51, 1–11.

Fournier GP, Gogarten JP (2010) Rooting the ribosomal tree of life. Molecular biology and evolution 27, 1792–801.

Gaby JC, Buckley DH (2014) A comprehensive aligned nifH gene database: a multipurpose tool for studies of nitrogen-fixing bacteria. Database□: the journal of biological databases and curation 2014, bau001.

Gallon JR (1981) The oxygen sensitivity of nitrogenase: a problem for biochemists and micro-organisms. Trends in Biochemical Sciences 6, 19–23.

Garcia AK, McShea H, Kolaczkowski B, Kaçar B (2020) Reconstructing the evolutionary history of nitrogenases: Evidence for ancestral molybdenum cofactor utilization. Geobiology gbi.12381.

Garvin J, Buick R, Anbar AD, Arnold GL, Kaufman AJ (2009a) Isotopic Evidence for an Aerobic Nitrogen Cycle in the Latest Archean. Science 323, 1045–1048.

Garvin J, Buick R, Anbar AD, Arnold GL, Kaufman AJ (2009b) Isotopic evidence for an aerobic nitrogen cycle in the latest Archean. Science 323, 1045–8.

Giblin A., Tobias CR, Song B, Weston N, Banta GT, Rivera-Monroy VH (2013) The importance of dissimilatory nitrate reduction to ammonium (DNRA) in the nitrogen cycle of coastal ecosystems. Oceanography 26, 124–131.

Gibson TM, Shih PM, Cumming VM, Fischer WW, Crockford PW, Hodskiss MSW, Worndle S, Creaser RA, Rainbird RH, Skulski TM, Halverson GP (2018) Precise age of Bangiomorpha pubescens dates the origin of eukaryotic photosynthesis. Geology 46, 135–138.

Godfrey L V., Falkowski PG (2009) The cycling and redox state of nitrogen in the Archaean ocean. Nature Geoscience 2, 725–729.

Gogarten JP, Townsend JP (2005) Horizontal gene transfer, genome innovation and evolution. Nature reviews. Microbiology 3, 679–87.

Hoffman PF, Abbot DS, Ashkenazy Y, Benn DI, Brocks JJ, Cohen PA, Cox GM, Creveling JR, Donnadieu Y, Erwin DH, Fairchild IJ, Ferreira D, Goodman JC, Halverson GP, Jansen MF, Hir G Le, Love GD, Macdonald FA, Maloof AC, Partin CA, Ramstein G, Rose BEJ, Rose C V., Sadler PM, Tziperman E, Voigt A, Warren SG (2017) Snowball Earth climate dynamics and Cryogenian geology-geobiology. Science Advances 3, e1600983.

Howard JB, Kechris KJ, Rees DC, Glazer AN (2013) Multiple amino acid sequence alignment nitrogenase component 1: insights into phylogenetics and structure-function relationships. PloS one 8, e72751.

Hug LA, Baker BJ, Anantharaman K, Brown CT, Probst AJ, Castelle CJ, Butterfield CN, Hernsdorf AW, Amano Y, Ise K, Suzuki Y, Dudek N, Relman DA, Finstad KM, Amundson R, Thomas BC, Banfield JF (2016) A new view of the tree of life. Nature Microbiology 1, 16048.

Hyatt D, Chen G-L, LoCascio PF, Land ML, Larimer FW, Hauser LJ (2010) Prodigal: prokaryotic gene recognition and translation initiation site identification. BMC Bioinformatics 11, 119.

Isson TT, Love GD, Dupont CL, Reinhard CT, Zumberge AJ, Asael D, Gueguen B, McCrow J, Gill BC, Owens J, Rainbird RH, Rooney AD, Zhao M-Y, Stueeken EE, Konhauser KO, John SG, Lyons TW, Planavsky NJ (2018) Tracking the rise of eukaryotes to ecological dominance with zinc isotopes. Geobiology 16, 341–352.

Joerger RD, Bishop PE, Evans HJ (1988) Bacterial Alternative Nitrogen Fixation Systems. CRC Critical Reviews in Microbiology 16, 1–14.

Johnson BW, Goldblatt C (2018) EarthN: A new earth system nitrogen model. Geochemistry, Geophysics, Geosystems.

Jones CM, Stres BB, Rosenquist M, Hallin S (2008) Phylogenetic analysis of nitrite, nitric oxide, and nitrous oxide respiratory enzymes reveal a complex evolutionary history for denitrification. Molecular Biology and Evolution 25, 1955–1966.

Kaiser JT, Hu Y, Wiig JA, Rees DC, Ribbe MW (2011) Structure of precursor bound NifEN: a nitrogenase FeMo cofactor maturase/insertase. Science (New York, N.Y.) 331, 91.

Kalyaanamoorthy S, Minh BQ, Wong TKF, Haeseler A von, Jermiin LS (2017) ModelFinder: fast model selection for accurate phylogenetic estimates. Nature Methods 14, 587–589.

Keable SM, Vertemara J, Zadvornyy OA, Eilers BJ, Danyal K, Rasmussen AJ, Gioia L De, Zampella G, Seefeldt LC, Peters JW (2018) Structural characterization of the nitrogenase molybdenum-iron protein with the substrate acetylene trapped near the active site. Journal of Inorganic Biochemistry 180, 129–134.

Kechris KJ, Lin JC, Bickel PJ, Glazer AN (2006) Quantitative exploration of the occurrence of lateral gene transfer by using nitrogen fixation genes as a case study. Proceedings of the National Academy of Sciences of the United States of America 103, 9584–9.

Kipp MAMA, Stüeken EEEE, Yun M, Bekker A, Buick R (2018) Pervasive aerobic nitrogen cycling in the surface ocean across the Paleoproterozoic Era. Earth and Planetary Science Letters 500, 117–126.

Kitts PA, Church DM, Thibaud-Nissen F, Choi J, Hem V, Sapojnikov V, Smith RG, Tatusova T, Xiang C, Zherikov A, DiCuccio M, Murphy TD, Pruitt KD, Kimchi A (2016) Assembly: a resource for assembled genomes at NCBI. Nucleic acids research 44, D73–80.

Knoll AH, Nowak MA (2017) The timetable of evolution. Science Advances 3, e1603076.

Koehler MC, Buick R, Barley ME (2019) Nitrogen isotope evidence for anoxic deep marine environments from the Mesoarchean Mosquito Creek Formation, Australia. Precambrian Research 320, 281–290.

Koehler MC, Buick R, Kipp MA, Stüeken EE, Zaloumis J (2018) Transient surface ocean oxygenation recorded in the ∼2.66-Ga Jeerinah Formation, Australia. Proceedings of the National Academy of Sciences of the United States of America 115, 7711–7716.

Koonin E V., Makarova KS, Aravind L (2001) Horizontal gene transfer in prokaryotes: Quantification and classification. Annual Review of Microbiology 55, 709–742.

Kozlov AM, Darriba D, Flouri T, Morel B, Stamatakis A (2019) RAxML-NG: a fast, scalable and user-friendly tool for maximum likelihood phylogenetic inference. Bioinformatics 35, 4453–4455.

Kuypers MMM, Marchant HK, Kartal B (2018) The microbial nitrogen-cycling network. Nature reviews. Microbiology 16, 263–276.

Lartillot N, Lepage T, Blanquart S (2009) PhyloBayes 3: A Bayesian software package for phylogenetic reconstruction and molecular dating. Bioinformatics 25, 2286–2288.

Lee CC, Hu Y, Ribbe MW (2010) Vanadium nitrogenase reduces CO. Science (New York, N.Y.) 329, 642.

Lepage T, Bryant D, Philippe H, Lartillot N (2007) A general comparison of relaxed molecular clock models. Molecular biology and evolution 24, 2669–80.

Lipschultz F, Wofsy SC, Ward BB, Codispoti LA, Friedrich G, Elkins JW (1990) Bacterial transformations of inorganic nitrogen in the oxygen-deficient waters of the Eastern Tropical South Pacific Ocean. Deep Sea Research Part A. Oceanographic Research Papers 37, 1513–1541.

Löscher CR, Kock A, Könneke M, LaRoche J, Bange HW, Schmitz RA (2012) Production of oceanic nitrous oxide by ammonia-oxidizing archaea. Biogeosciences 9, 2419–2429.

Luo G, Junium CK, Izon G, Ono S, Beukes NJ, Algeo TJ, Cui Y, Xie S, Summons RE (2018) Nitrogen fixation sustained productivity in the wake of the Palaeoproterozoic Great Oxygenation Event. Nature Communications 9, 978.

Lyons TW, Reinhard CT, Planavsky NJ (2014) The rise of oxygen in Earth’s early ocean and atmosphere. Nature 506, 307–315.

Magnabosco C, Moore KR, Wolfe JM, Fournier GP (2018) Dating phototrophic microbial lineages with reticulate gene histories. Geobiology 16, 179–189.

Marty B, Zimmermann L, Pujol M, Burgess R, Philippot P (2013) Nitrogen isotopic composition and density of the Archean atmosphere. Science 342, 101–104.

Mather TA, Pyle DM, Allen AG (2004) Volcanic source for fixed nitrogen in the early Earth’s atmosphere. Geology 32, 905.

McGlynn SE, Boyd ES, Peters JW, Orphan VJ (2012) Classifying the metal dependence of uncharacterized nitrogenases. Frontiers in microbiology 3, 419.

McInerney JO, McNally A, O’Connell MJ (2017) Why prokaryotes have pangenomes. Nature Microbiology 2, 17040.

Mendler K, Chen H, Parks DH, Lobb B, Hug LA, Doxey AC (2019) AnnoTree: visualization and exploration of a functionally annotated microbial tree of life. Nucleic Acids Research 47, 4442–4448.

Michiels CC, Darchambeau F, Roland FAE, Morana C, Llirós M, García-Armisen T, Thamdrup B, Borges A V., Canfield DE, Servais P, Descy J-P, Crowe SA (2017) Iron-dependent nitrogen cycling in a ferruginous lake and the nutrient status of Proterozoic oceans. Nature Geoscience 10, 217–221.

Miller RW, Eady RR (1988) Molybdenum and vanadium nitrogenases of Azotobacter chroococcum. Low temperature favours N2 reduction by vanadium nitrogenase. Biochemical Journal 256, 429–432.

Mirkin BG, Fenner TI, Galperin MY, Koonin E V (2003) Algorithms for computing parsimonious evolutionary scenarios for genome evolution, the last universal common ancestor and dominance of horizontal gene transfer in the evolution of prokaryotes. BMC Evolutionary Biology 3, 2.

Mojzsis SJ, Arrhenius G, McKeegan KD, Harrison TM, Nutman AP, Friend CRL (1996) Evidence for life on Earth before 3,800 million years ago. Nature 384, 55–59.

Moore EK, Hao J, Spielman SJ, Yee N (2020) The evolving redox chemistry and bioavailability of vanadium in deep time. Geobiology 18, 127–138.

Moore EK, Jelen BI, Giovannelli D, Raanan H, Falkowski PG (2017) Metal availability and the expanding network of microbial metabolisms in the Archaean eon. Nature Geoscience 10, 629–636.

Moulana A, Anderson RE, Fortunato CS, Huber JA (2020) Selection is a significant driver of gene gain and loss in the pangenome of the bacterial genus Sulfurovum in geographically distinct deep-sea hydrothermal vents. mSystems.

Mus F, Alleman AB, Pence N, Seefeldt LC, Peters JW (2018) Exploring the alternatives of biological nitrogen fixation. Metallomics 10, 523–538.

Mus F, Colman DR, Peters JW, Boyd ES (2019) Geobiological feedbacks, oxygen, and the evolution of nitrogenase. Free Radical Biology and Medicine.

Navarro-González R, McKay CP, Mvondo DN (2001) A possible nitrogen crisis for Archaean life due to reduced nitrogen fixation by lightning. Nature 412, 61–64.

Noffke N, Beukes N, Bower D, Hazen RM, Swift DJP (2007) An actualistic perspective into Archean worlds -(cyano-)bacterially induced sedimentary structures in the siliciclastic Nhlazatse Section, 2.9 Ga Pongola Supergroup, South Africa. Geobiology 6, 5–20.

Nutman AP, Bennett VC, Friend CRL, Kranendonk MJ Van, Chivas AR (2016) Rapid emergence of life shown by discovery of 3,700-million-year-old microbial structures. Nature 537, 535–538.

Ogata H, Goto S, Sato K, Fujibuchi W, Bono H, Kanehisa M (1999) KEGG: Kyoto Encyclopedia of Genes and Genomes. Nucleic Acids Research 27, 29–34.

Olson SL, Kump LR, Kasting JF (2013) Quantifying the areal extent and dissolved oxygen concentrations of Archean oxygen oases. Chemical Geology 362, 35–43.

Ossa Ossa F, Hofmann A, Spangenberg JE, Poulton SW, Stüeken EE, Schoenberg R, Eickmann B, Wille M, Butler M, Bekker A (2019) Limited oxygen production in the Mesoarchean ocean. Proceedings of the National Academy of Sciences of the United States of America 116, 6647–6652.

Pang K, Tang Q, Chen L, Wan B, Niu C, Yuan X, Xiao S (2018) Nitrogen-Fixing Heterocystous Cyanobacteria in the Tonian Period. Current Biology 28, 616–622.

Pang K, Tang Q, Schiffbauer JD, Yao J, Yuan X, Wan B, Chen L, Ou Z, Xiao S (2013) The nature and origin of nucleus-like intracellular inclusions in Paleoproterozoic eukaryote microfossils. Geobiology 11, 499–510.

Planavsky NJ, Asael D, Hofmann A, Reinhard CT, Lalonde S V, Knudsen A, Wang X, Ossa Ossa F, Pecoits E, Smith AJB, Beukes NJ, Bekker A, Johnson TM, Konhauser KO, Lyons TW, Rouxel OJ (2014) Evidence for oxygenic photosynthesis half a billion years before the Great Oxidation Event. Nature Geoscience 7, 283–286.

Polz MF, Alm EJ, Hanage WP (2013) Horizontal gene transfer and the evolution of bacterial and archaeal population structure. Trends in Genetics 29, 170–5.

Popa O, Hazkani-Covo E, Landan G, Martin W, Dagan T (2011) Directed networks reveal genomic barriers and DNA repair bypasses to lateral gene transfer among prokaryotes. Genome research 21, 599–609.

Raymond J, Siefert JL, Staples CR, Blankenship RE (2004) The natural history of nitrogen fixation. Molecular Biology and Evolution 21, 541–54.

Reinhard CT, Planavsky NJ, Robbins LJ, Partin CA, Gill BC, Lalonde S V., Bekker A, Konhauser KO, Lyons TW (2013) Proterozoic ocean redox and biogeochemical stasis. Proceedings of the National Academy of Sciences 110, 5357–5362.

Ren M, Feng X, Huang Y, Wang H, Hu Z, Clingenpeel S, Swan BK, Fonseca MM, Posada D, Stepanauskas R, Hollibaugh JT, Foster PG, Woyke T, Luo H (2019) Phylogenomics suggests oxygen availability as a driving force in Thaumarchaeota evolution. The ISME Journal 13, 2150–2161.

Revell LJ (2012) phytools: an R package for phylogenetic comparative biology (and other things). Methods in Ecology and Evolution 3, 217–223.

Roberson AL, Roadt J, Halevy I, Kasting JF (2011) Greenhouse warming by nitrous oxide and methane in the Proterozoic Eon. Geobiology 9, 313–320.

Rosing MT (1999) 13C-Depleted carbon microparticles in &gt;3700-Ma sea-floor sedimentary rocks from west greenland. Science (New York, N.Y.) 283, 674–6.

Sánchez-Baracaldo P, Ridgwell A, Raven JA (2014) A neoproterozoic transition in the marine nitrogen cycle. Current Biology 24, 652–657.

Santos PC Dos, Fang Z, Mason SW, Setubal JC, Dixon R (2012) Distribution of nitrogen fixation and nitrogenase-like sequences amongst microbial genomes. BMC Genomics 13, 162.

Schidlowski M (1988) A 3,800-million-year isotopic record of life from carbon in sedimentary rocks. Nature 333, 313–318.

Schidlowski M, Hayes JM, Kaplan IR (1983) Isotopic inferences of ancient biochemistries: Carbon, sulfur, hydrogen, and nitrogen. In: Earth’s Earliest Biosphere, Its Origin and Evolution (ed. Schopf JW). Princeton University Press, Princeton, N.J., pp. 149–186.

Schoepp-Cothenet B, Lis R van, Atteia A, Baymann F, Capowiez L, Ducluzeau A-L, Duval S, Brink F ten, Russell MJ, Nitschke W (2013) On the universal core of bioenergetics. Biochimica et Biophysica Acta (BBA) - Bioenergetics 1827, 79–93.

Schwartz JH, Maresca B (2006) Do Molecular Clocks Run at All? A Critique of Molecular Systematics. Biological Theory 1, 357–371.

Scott C, Lyons TW, Bekker A, Shen Y, Poulton SW, Chu X, Anbar AD (2008) Tracing the stepwise oxygenation of the Proterozoic ocean. Nature 452, 456–459.

Shang M, Tang D, Shi X, Zhou L, Zhou X, Song H, Jiang G (2019) A pulse of oxygen increase in the early Mesoproterozoic ocean at ca. 1.57–1.56 Ga. Earth and Planetary Science Letters 527, 115797.

Som SM, Buick R, Hagadorn JW, Blake TS, Perreault JM, Harnmeijer JP, Catling DC (2016) Earth’s air pressure 2.7 billion years ago constrained to less than half of modern levels. Nature Geoscience 9, 448–451.

Stamatakis A (2014) RAxML version 8: a tool for phylogenetic analysis and post-analysis of large phylogenies. Bioinformatics 30, 1312–1313.

Stanton CL, Reinhard CT, Kasting JF, Ostrom NE, Haslun JA, Lyons TW, Glass JB (2018) Nitrous oxide from chemodenitrificationLJ: A possible missing link in the Proterozoic greenhouse and the evolution of aerobic respiration 1–13.

Stein LY, Klotz MG (2016) The nitrogen cycle. Current Biology 26, R94–R98.

Stolz JF, Basu P (2002) Evolution of Nitrate Reductase: Molecular and Structural Variations on a Common Function. ChemBioChem 3, 198–206.

Stüeken EE, Buick R, Guy BM, Koehler MC (2015) Isotopic evidence for biological nitrogen fixation by molybdenum-nitrogenase from 3.2 Gyr. Nature.

Stüeken EE, Catling DC, Buick R (2012) Contributions to late Archaean sulphur cycling by life on land. Nature Geoscience 5, 722–725.

Stüeken EE, Kipp MA, Koehler MC, Buick R (2016a) The evolution of Earth’s biogeochemical nitrogen cycle. Earth-Science Reviews 160, 220–239.

Stüeken EE, Kipp MA, Koehler MC, Schwieterman EW, Johnson B, Buick R (2016b) Modeling *p* N _2_ through Geological Time: Implications for Planetary Climates and Atmospheric Biosignatures. Astrobiology 16, 949–963.

Summers DP, Chang S (1993) Prebiotic ammonia from reduction of nitrite by iron (II) on the early Earth. Nature 365, 630–633.

The Uniprot Consortium (2017) UniProt: the universal protein knowledgebase. Nucleic Acids Research 45, D158–D169.

Voss M, Bange HW, Dippner JW, Middelburg JJ, Montoya JP, Ward B (2013) The marine nitrogen cycle: recent discoveries, uncertainties and the potential relevance of climate change. Philosophical Transactions of the Royal Society B: Biological Sciences 368, 20130121–20130121.

Wada E, Hatton A (1971) Nitrite metabolism in the euphotic layer of the central North Pacific Ocean. Limnology and Oceanography 16, 766–772.

Waterhouse AM, Procter JB, Martin DMA, Clamp M, Barton GJ (2009) Jalview Version 2--a multiple sequence alignment editor and analysis workbench. Bioinformatics 25, 1189–1191.

Weiss MC, Sousa FL, Mrnjavac N, Neukirchen S, Roettger M, Nelson-Sathi S, Martin WF (2016) The physiology and habitat of the last universal common ancestor. Nature Microbiology 1, 16116.

Yang J, Junium CK, Grassineau N V., Nisbet EG, Izon G, Mettam C, Martin A, Zerkle AL (2019) Ammonium availability in the Late Archaean nitrogen cycle. Nature Geoscience 12, 553–557.

Zaremba-Niedzwiedzka K, Caceres EF, Saw JH, Bäckström D, Juzokaite L, Vancaester E, Seitz KW, Anantharaman K, Starnawski P, Kjeldsen KU, Stott MB, Nunoura T, Banfield JF, Schramm A, Baker BJ, Spang A, Ettema TJG (2017) Asgard archaea illuminate the origin of eukaryotic cellular complexity. Nature 541, 353–358.

Zerkle AL, Mikhail S (2017) The geobiological nitrogen cycle: From microbes to the mantle. Geobiology.

Zerkle AL, Poulton SW, Newton RJ, Mettam C, Claire MW, Bekker A, Junium CK(2017a) Onset of the aerobic nitrogen cycle during the Great Oxidation Event.Nature 542, 465–467.

Zerkle AL, Poulton SW, Newton RJ, Mettam C, Claire MW, Bekker A, Junium CK(2017b) Onset of the aerobic nitrogen cycle during the Great Oxidation Event.Nature 542, 465–467.

Zhang K, Zhu X, Wood RA, Shi Y, Gao Z, Poulton SW (2018) Oxygenation of the Mesoproterozoic ocean and the evolution of complex eukaryotes. Nature Geoscience 11, 345–350.

