## Supplementary tables for "Radiation of nitrogen-metabolizing enzymes across the tree of life tracks environmental transitions in Earth history"

**All Newick files, chronograms and genome lists have been deposited to FigShare at** <https://figshare.com/projects/Radiation_of_nitrogen_cycling_genes_across_the_tree_of_life/87461>**.**

| **Calibration Event** | **Liberal (Ga)** | **Refs** |
| --- | --- | --- |
| LUCA (set as root prior) | 4100, S.D. 200 | (Bell *et al.*, 2015) |
| Origin of Cyanobacteria | >3200 | (Betts *et al.*, 2018) |
| Origin of Methanogenesis | >3510 | (Wolfe & Fournier, 2018) |
| Origin of Eukaryotes | >3200 | (Javaux *et al.*, 2010) |
| Origin of plastids/Rhodophytes diverge | >1200 | (Butterfield, 2000) |
| Akinetes diverge from cyanobacteria lacking cell differentiation | >1500 | (Golubic *et al.*, 1995) |

**Supplementary Table 1.** Fossil calibration points used in Phylobayes runs. Calibration points were set as the hard constraint indicating the latest point by which something arose. These liberal time points reflect the earliest dates for which there is evidence of these events based on the current scientific literature. The liberal calibration points were used comparison for the generation of chronograms, but the resulting chronograms were not used for subsequent analysis.

| **Gene** | **Upper node date** | **95% confidence interval** | **Lower node date** | **95% confidence interval** | **Midpoint between nodes** | **Geologic era** |
| --- | --- | --- | --- | --- | --- | --- |
| **vnfD** | 819.68 | 532.685, 1179.96 | 241.90 | 138.984, 409.921 | 530.79 | **Neoproterozoic** |
| **vnfK** | 819.68 | 532.685, 1179.96 | 241.90 | 138.984, 409.921 | 530.79 | **Neoproterozoic** |
| **nosZ** | 1034.26 | 800.348, 1314.43 | 790.53 | 560.97, 1066.4 | 912.40 | **Neoproterozoic** |
| **nirK** | 1161.98 | 913.226, 1457.9 | 1004.18 | 774.694, 1278.81 | 1083.08 | **Mesoproterozoic** |
| **nirS** | 1717.48 | 1422.29, 2051.72 | 1165.82 | 918.685, 1463.03 | 1441.65 | **Mesoproterozoic** |
| **norB** | 1717.48 | 1422.29, 2051.72 | 1165.82 | 918.685, 1463.03 | 1441.65 | **Mesoproterozoic** |
| **anfD** | 2190.75 | 1847.38, 2541.5 | 1585.61 | 1291.61, 1916.62 | 1888.18 | **Paleoproterozoic** |
| **anfK** | 2190.75 | 1847.38, 2541.5 | 1585.61 | 1291.61, 1916.62 | 1888.18 | **Paleoproterozoic** |
| **nasA** | 2583.62 | 2281.26, 2904 | 1967.99 | 1644.3, 2318.01 | 2275.81 | **Paleoproterozoic** |
| **nirB** | 2776.44 | 2353.91, 3166.91 | 2355.87 | 1882.42, 2804.97 | 2566.15 | **Neoarchean** |
| **nifK** | 3372.12 | 3081.12, 3679.08 | 2601.08 | 2042.03, 3092.06 | 2986.60 | **Mesoarchean** |
| **nrfA** | 3138.86 | 2856.06, 3439.62 | 2946.21 | 2608.87, 3270.37 | 3042.54 | **Mesoarchean** |
| **narG** | 3138.86 | 2856.06, 3439.62 | 3061.22 | 2773.92, 3367.72 | 3100.04 | **Mesoarchean** |
| **nxrA** | 3138.86 | 2856.06, 3439.62 | 3061.22 | 2773.92, 3367.72 | 3100.04 | **Mesoarchean** |
| **napA** | 3256.96 | 2979.4, 3558.14 | 3138.86 | 2856.06, 3439.62 | 3197.91 | **Mesoarchean** |
| **nifH** | 3256.96 | 2979.4, 3558.14 | 3138.86 | 2856.06, 3439.62 | 3197.91 | **Mesoarchean** |
| **nifD** | 3406.74 | 3138.44, 3704.79 | 3256.96 | 2979.4, 3558.14 | 3331.85 | **Paleoarchean** |

**Supplementary Table 2**. Inferred birth dates for nitrogen-cycling genes based on the chronogram generated using the UGAM clock model with conservative calibration points. All gene birth events are inferred to occur between nodes on the species chronogram, and therefore our methods do not allow us to infer specific dates for gene birth events. Here we list the inferred timing for the earliest possible timing (upper node) and latest possible timing (lower node) with 95% confidence intervals as reported by PhyloBayes, as well as the calculated midpoint between the two node times. Times are reported in millions of years ago. The geological era reported in the right column corresponds to the midpoint between the upper and lower node. Given the uncertainty inherent in the estimation of node timings and inference in events, these are provided for geological reference and should not be taken as definitive timings. The results in this table are equivalent to the data presented in Table 4, but were generated using the UGAM clock model rather than the CIR clock model.
