## Supplementary figures for "Radiation of nitrogen-metabolizing enzymes across the tree of life tracks environmental transitions in Earth history"

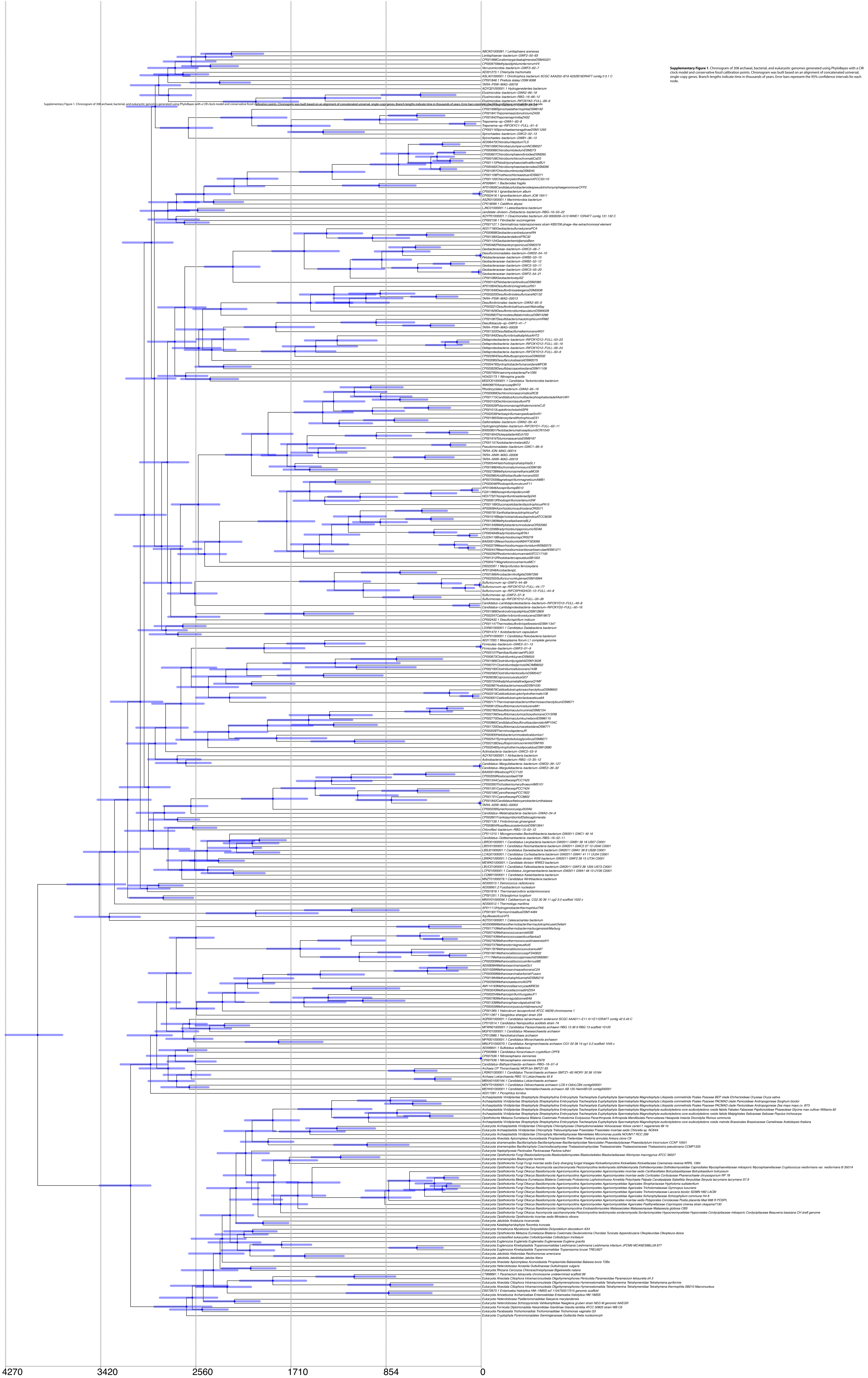


**Supplementary Figure 1**. Chronogram of 308 archaeal, bacterial, and eukaryotic genomes generated using PhyloBayes with a CIR clock model and conservative fossil calibration points. Chronogram was built based on an alignment of concatenated universal, single-copy genes. Branch lengths indicate time in thousands of years. Error bars represent the 95% confidence intervals for each node.

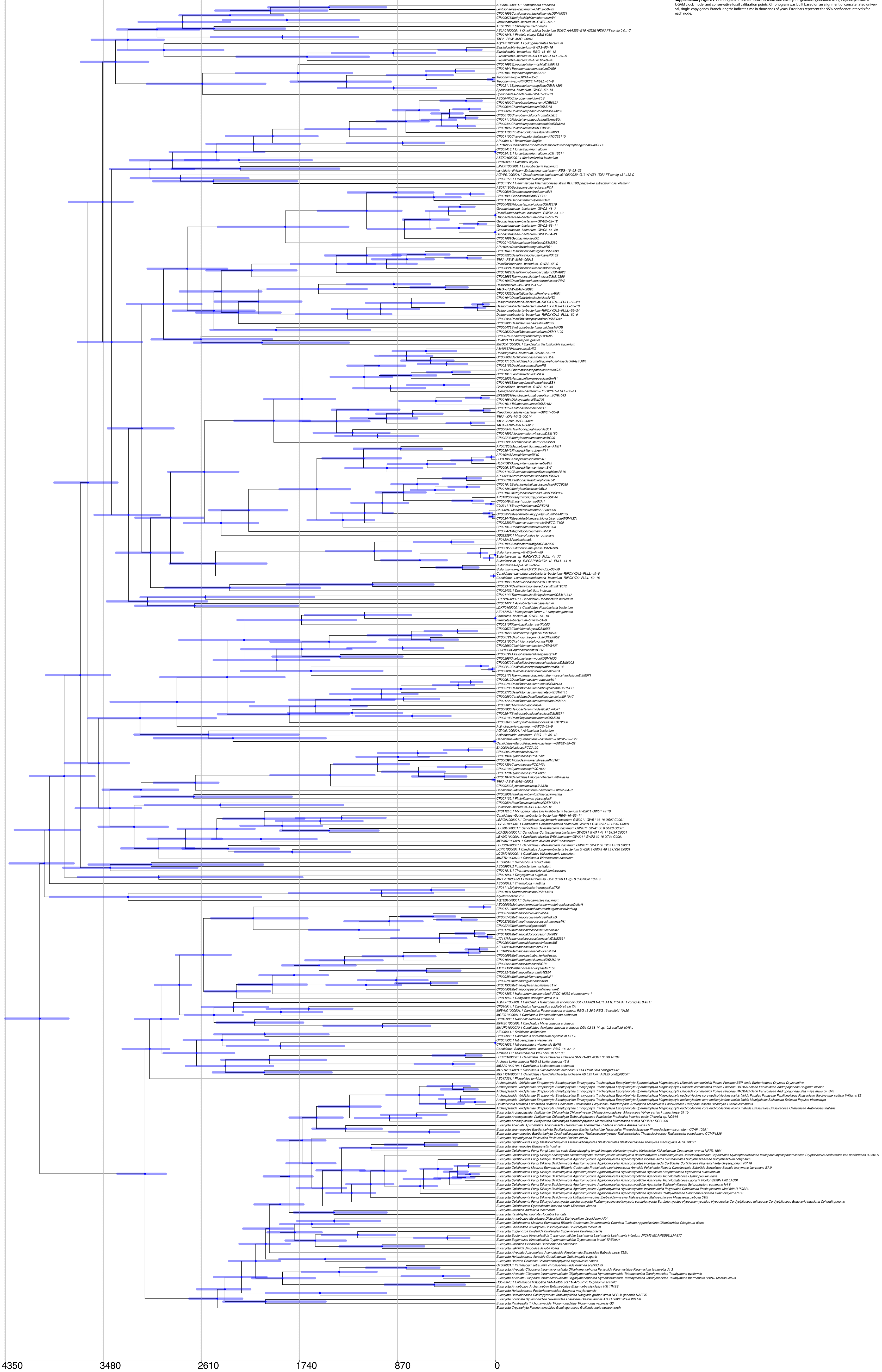


**Supplementary Figure 2**. Chronogram of 308 archaeal, bacterial, and eukaryotic genomes generated using PhyloBayes with a UGAM clock model and conservative fossil calibration points. Chronogram was built based on an alignment of concatenated universal, single-copy genes. Branch lengths indicate time in thousands of years. Error bars represent the 95% confidence intervals for each node.

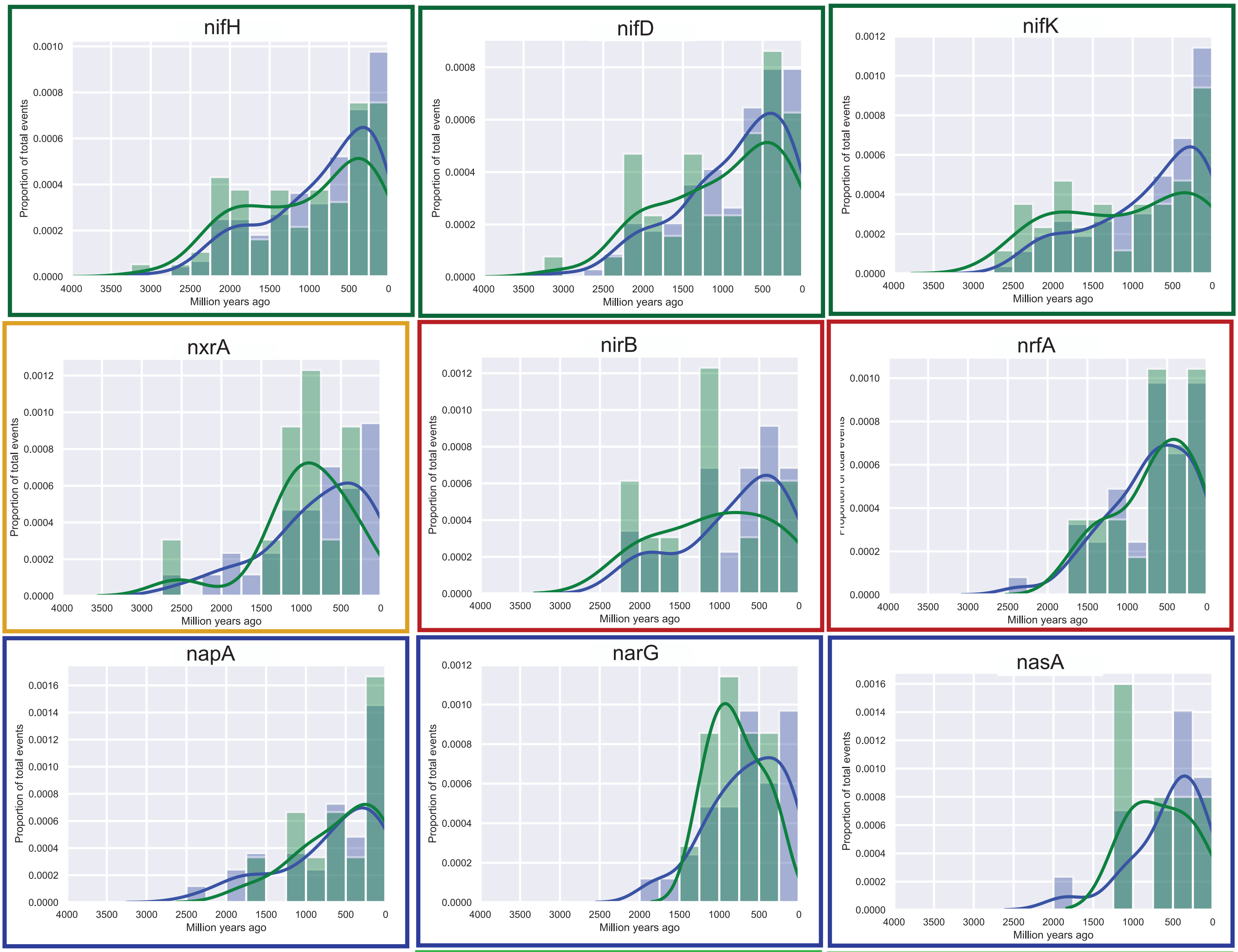


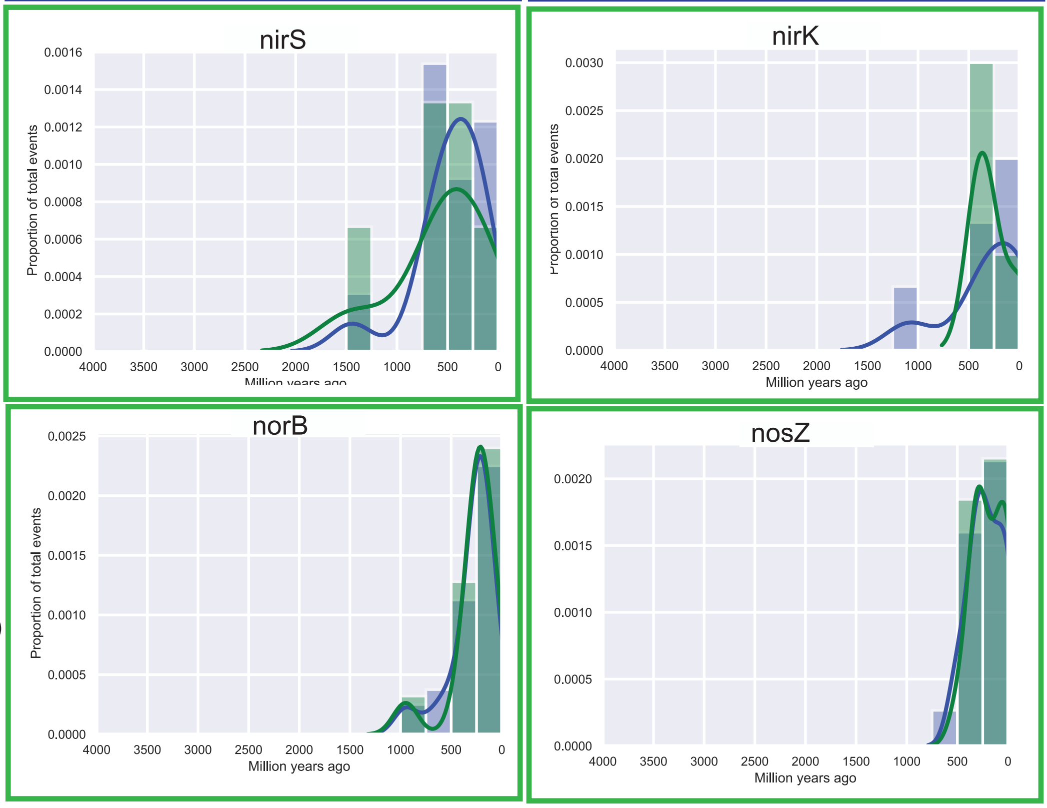


**Supplementary Figure 3.** Histograms and density plots of all speciation, loss, duplication and horizontal gene transfer (HGT) events. All events are shown in blue, HGT events alone are shown in green. Events were inferred by reconciling phylogenetic trees of nitrogen-cycling genes with a chronogram generated using a UGAM clock model and conservative calibration points. Y-axis represents proportion of all events (blue) or HGT events only (green). Borders around graphs indicate the metabolic process that the gene is involved in. Histograms are ordered according to the earliest time point at which a speciation, loss, duplication, or HGT event occurred.

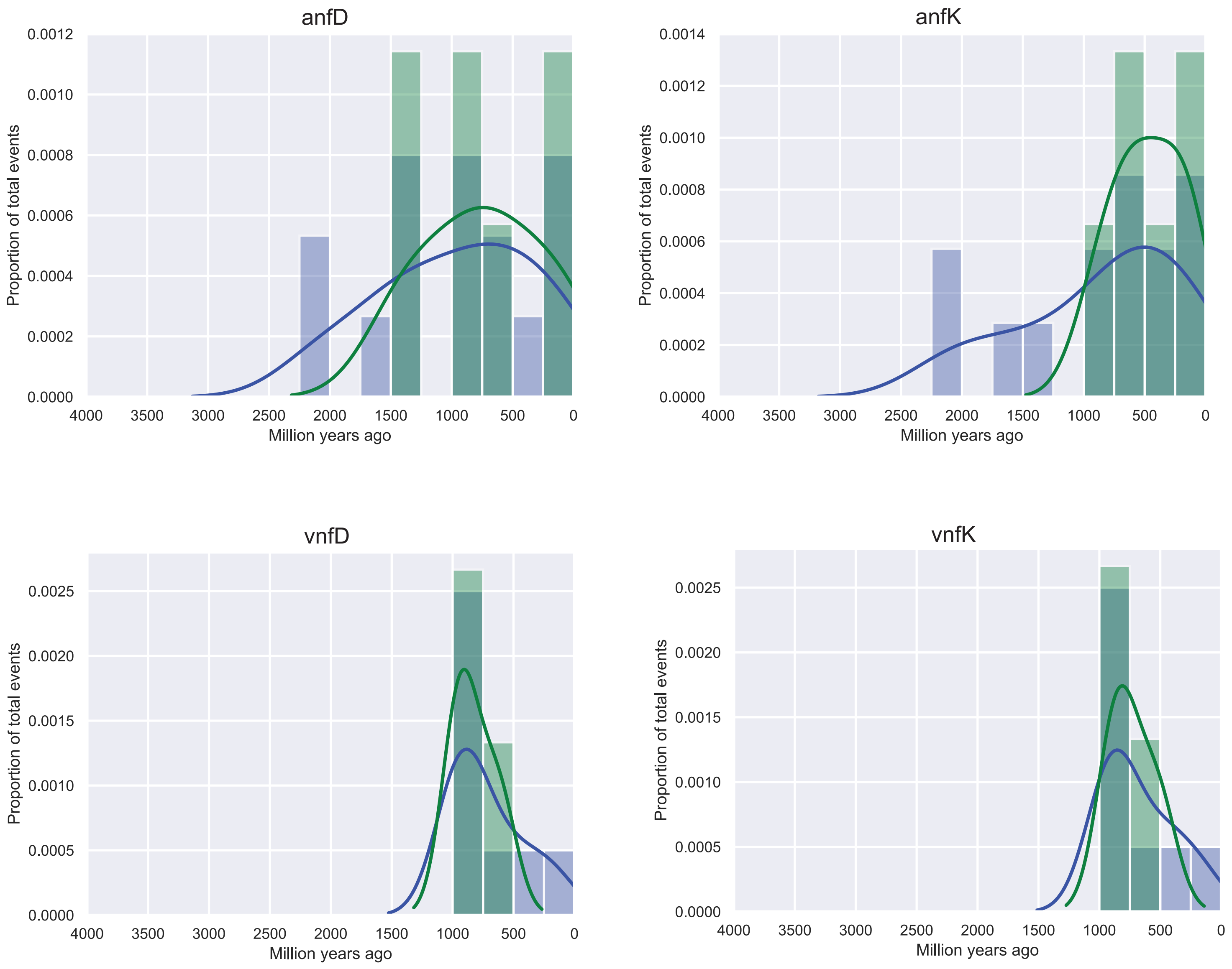


**Supplementary Figure 4**. Histograms and density plots of all speciation, loss, duplication, and HGT events (blue) and just horizontal gene transfer events (green) for anf and vnf genes inferred from the species chronogram generated using the CIR clock model with conservative calibration points.

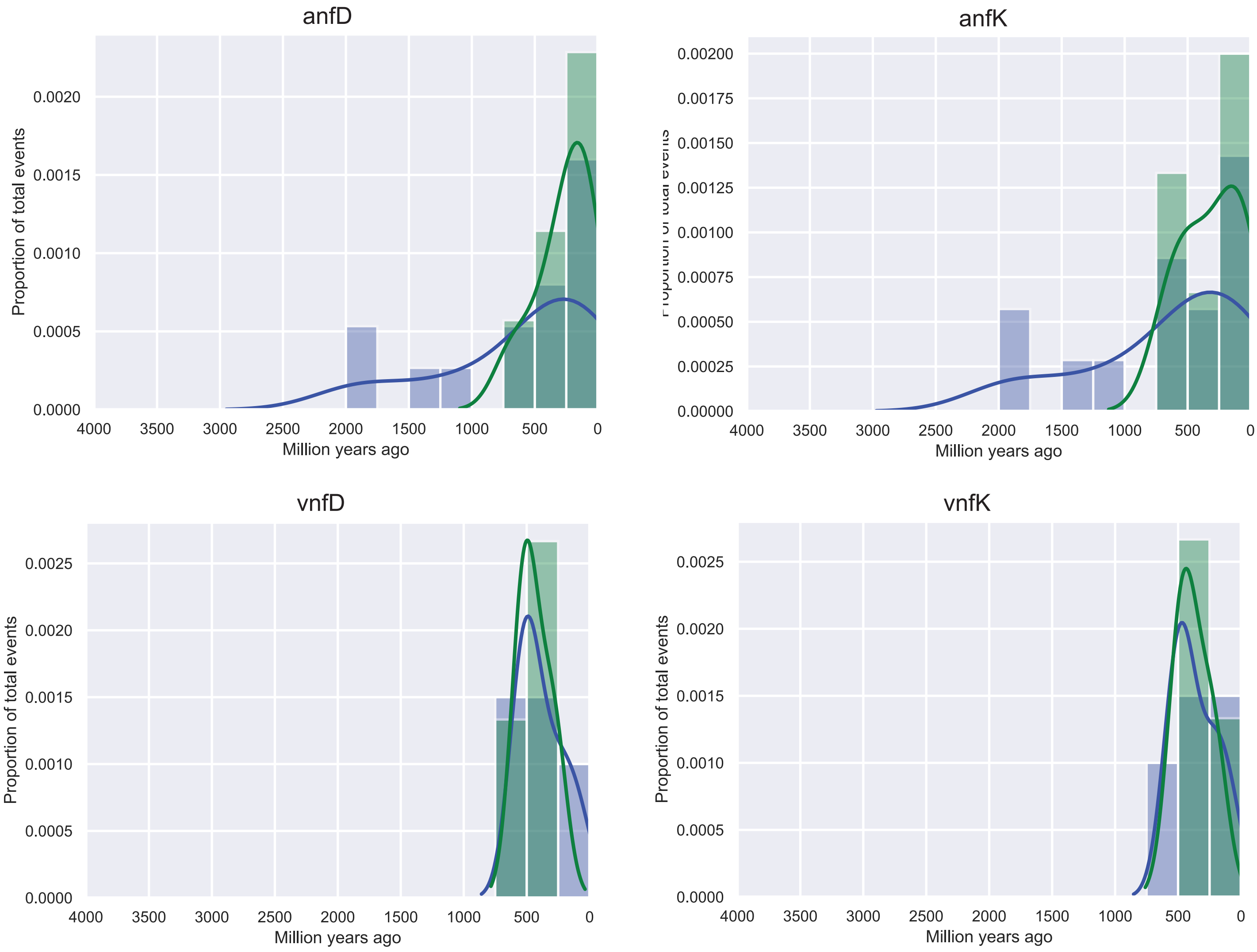


**Supplementary Figure 5.** Histograms and density plots of all speciation, loss, duplication, and HGT events (blue) and just horizontal gene transfer events (green) for anf and vnf genes inferred from the species chronogram generated using the UGAM clock model with conservative calibration points.
